# Numerous post-translational modifications of RNA polymerase II subunit Rpb4 link transcription to post-transcriptional mechanisms

**DOI:** 10.1101/798603

**Authors:** Stephen Richard, Lital Gross, Jonathan Fischer, Keren Bendalak, Tamar Ziv, Shira Urim, Mordechai Choder

## Abstract

Rpb4/7 binds RNA Polymerase II (Pol II) transcripts co-transcriptionally and accompanies them throughout their lives. By virtue of its capacity to interact with key regulators (e.g., Pol II, eIF3, Pat1) both temporarily and spatially, Rpb4/7 regulates the major stages of the mRNA lifecycle. Here we show that Rpb4/7 can undergo over 100 combinations of post-translational modifications (PTMs). Remarkably, the Rpb4/7 PTMs repertoire changes as the mRNA/Rpb4/7 complex progresses from one stage to the next. A mutagenesis approach in residues that undergo PTMs suggests that temporal Rpb4 PTMs regulate its interactions with key regulators of gene expression that control transcriptional and post-transcriptional stages. Moreover, one mutant type specifically affects mRNA synthesis despite its normal association with Pol II, whereas the other affects both mRNA synthesis and decay; both types disrupt the balance between mRNA synthesis and decay (‘mRNA buffering’) and the cell’s capacity to respond to the environment. Taken together, we propose that temporal Rpb4/7 PTMs are involved in cross talks among the various stages of the mRNA lifecycle.

## Introduction

Post-translational modifications (PTMs) play fundamental roles in regulating proteins function in a wide variety of cellular processes. PTMs can diversify a protein’s function, modulate its interaction with other proteins and dynamically coordinate its involvement in signaling networks. PTMs are involved in regulating almost all cellular events, including gene expression, signal transduction, protein-protein interaction, cell-cell interaction, and communication between the intracellular and extracellular environment (Deribe et al., 2010). PTMs play a part in coordinating the co-transcriptional assembly, as well as remodeling and export of mRNP (Tutucci and Stutz, 2011). Nonetheless, relatively little is known about their impact on the linkage between distinct stages of gene expression (e.g., mRNA synthesis and decay), or on cross talks between distinct machineries or functional complexes.

The yeast *S. cerevisiae* Rpb4 and Rpb7 function as a heterodimer that mediates transcription initiation, elongation and polyadenylation as well as dephosphorylation of the C-terminal domain of Rpb1 and gene looping (Allepuz-Fuster et al., 2014; Babbarwal et al., 2014; Choder and Young, 1993; Orlicky et al., 2001; Rosenheck and Choder, 1998; Verma-Gaur et al., 2008)(Runner et al., 2008). Rpb4/7 binds Pol II transcripts co-transcriptionally and then accompanies the mRNA throughout its life. By binding key regulatory factors, such as components of the translation initiation factor 3 Nip1 and Hcr1 (Harel-Sharvit et al., 2010; Villanyi et al., 2014), or components of the decay complex, consisting of Pat1, Lsm2, and Not5 (Lotan et al., 2007, 2005; Villanyi et al., 2014), Rpb4/7 modulates mRNA export (Farago et al., 2003), translation (Harel-Sharvit et al., 2010; Villanyi et al., 2014) and decay (Bloom et al., 2013; Goler-Baron et al., 2008; Lotan et al., 2007, 2005). The mechanism underlying Rpb4/7’s capacity to bind these factors spatially and temporarily is unknown.

The role of Rpb4 in the linkage between mRNA synthesis and decay is controversial. Some investigators proposed that mRNA decay rate is linked to transcription rate by default and that Rpb4 impact on mRNA decay is merely a consequence of its effect on transcription (Schulz et al., 2014; Sun et al., 2012) while others proposed that the linkage is a regulated process involving many factors (Haimovich et al., 2013b, 2013a; Timmers and Tora, 2018). The latter proposals posit that Rpb4 plays two distinct roles in mRNA synthesis and decay (Duek et al., 2018; Goler-Baron et al., 2008; Lotan et al., 2007, 2005).

Here we show that transient PTMs affect the capacity of Rpb4/7 to temporarily bind a number of proteins and to regulate mRNA buffering by linking transcription and mRNA decay.

## Results

### Numerous isoforms of Rpb4 and Rpb7 are found by two-dimensional gel electrophoresis

Western analyses of tandem affinity purified Rpb4 suggested that it is composed of more than one isoform that migrated differently on SDS PAGE (Fig. 1A). The multiple functions assigned to Rpb4/7 (see Introduction) and the high MW bands, which were detected even following purification in the presence of a strong denaturation agent guanidine hydrochloride (Fig. 1B), encouraged us to examine the purified Rpb4 and Rpb7 by two dimensional gel electrophoresis (2D), composed of isoelectric focusing dimension followed by standard SDS-polyacrylamide gel. In several biological replicates, we have reproducibly observed at least 14 major Rpb4 isoforms (usually 20) (Fig. 1C), and many other minor isoforms (Fig. 1D). The high MW species are also detected as a number of isoforms (Fig. 1D). We also stained one of our 2D gels with Coomassie blue and cut out a few spots, including the 50 Kd spots, and analyzed them by mass spectrometry (MS). They all contained Rpb4 peptides (results not shown). We also probed these membranes with anti-Rpb7 and observed ~ 5 isoforms (Fig. 1E). Since Rpb4 is more heavily modified, we focused on the PTMs of this protein.

**Fig. 1.**
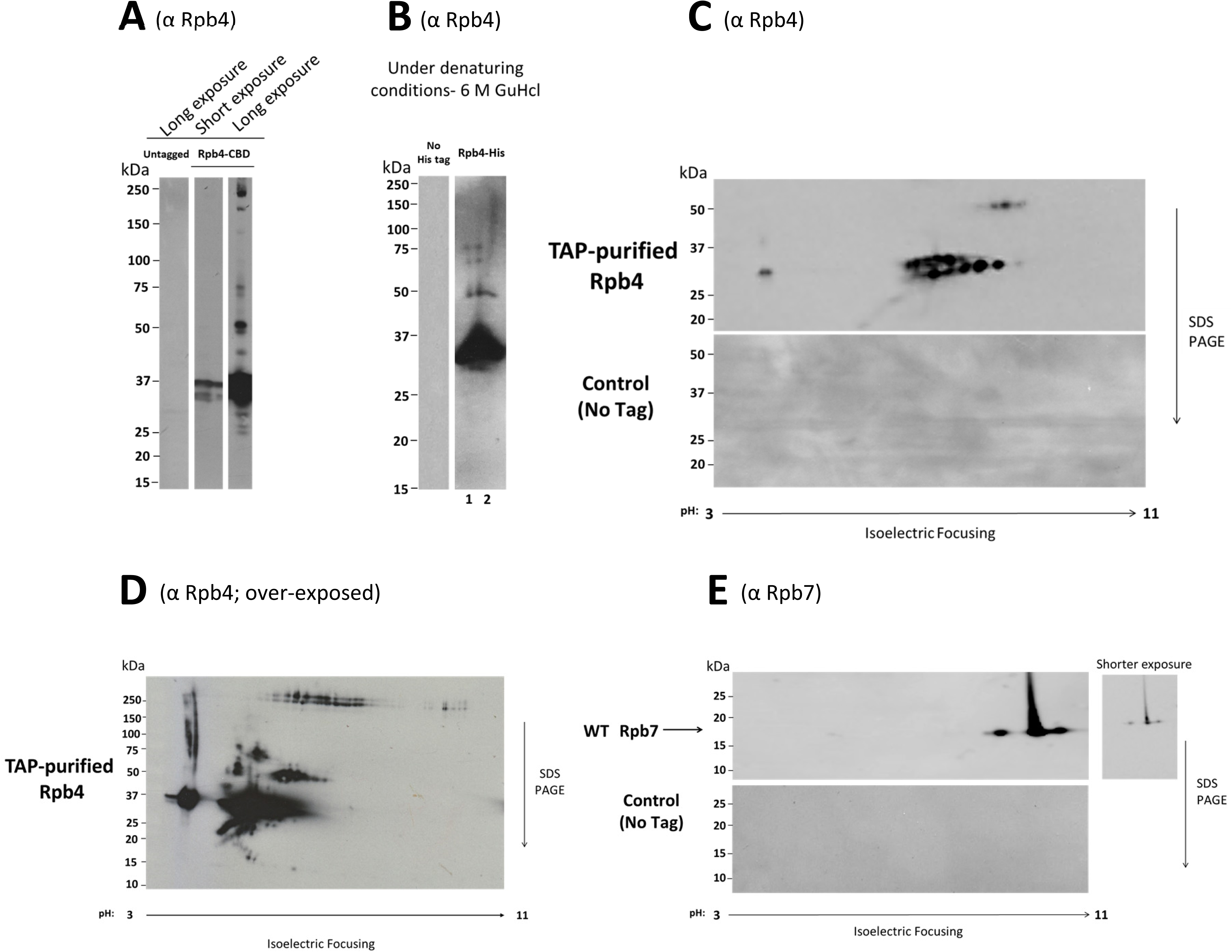
Rpb4 is separated by 2D gel into several isoforms. (A) One dimensional SDS PAGE reveals several Rpb4 isoforms. Rpb4 was tandemly affinity purified (Gavin et al., 2002) from *S. cerevisiae* (ySR007) extract and analyzed by a standard western blot analysis using anti-Rpb4 antibodies. (B) His-TEV-Protein A (HTP)-tagged Rpb4 was tandemly affinity-purified from ySR057 cell extract, using IgG resin as the first step followed by Ni-NTA agarose resin in the presence of 6M guanidine hydro-chloride (a strong detergent) as the second step (Van Nues et al., 2017). The purified protein was analyzed as in A. (C) Tandemly affinity purified Rpb4 was resolved by a two-dimensional gel electrophoresis (2D) (see Methods) followed by western analysis with anti-Rpb4 antibodies. (D) Overexposed membrane shown in C. (E) Rpb7 isoforms. The same membrane was reacted with anti-Rpb7 Abs.

### Mass Spectrometry analysis of Rpb4 reveals various kinds of PTMs

The isoform profile of Rpb4 strongly suggests that Rpb4 is post-translationally modified. Tandem affinity purified Rpb4 was analyzed by Mass Spectrometry (MS) for Post Translational Modifications (PTMs). In at least three biological replicates, we found about 37 modifications, summarized in Fig. S1A. Examples of annotated MS/MS spectra of modified peptides are shown in Fig. S1B and C. We detected peptides with two additional glycine residues (Gly Gly-Lys peptide) (Fig. S1A, and C), suggesting the addition of either ubiquitin or neddyl (Wagner et al., 2011). However, despite our persistent efforts, we could not detect any of these additions by MS or by specific antibodies. Moreover, we could not detect any ubiquitin ligase or ubiquitin-binding protein in our MS data, nor could we detect high MW bands in the 2D profiles that were sensitive to K to R replacement. It is possible that ubiquitylation/neddylation is transient and short lived, or that the Gly-Gly additions represent a new type of modification. We found methylations in a row of four glutamic acid residues (Fig. S2A), two of which are conserved from yeast to human (Fig. S2B), which are present in a loop which protrudes from the main Rpb4/7 structure (Fig. S2C-D). Methylation of glutamic acid is a relatively unexplored modification that has been reported to function in sensing and transducing chemical signals in *E.coli* by modulating interaction between protein domains (Clarke, 1993; Stock et al., 1992). These residues are named herein “E19-22 motif”. We were intrigued by its location in Rpb4/7 structure (Fig. S2C) and in the context of Pol II (Fig. S2E) and decided to investigate it further.

### Rpb4 PTMs respond to the environment and mutations in modified residues affect cell proliferation at 37°C

Rpb4 function becomes critical at temperatures above 32°C (Choder and Young, 1993; Farago et al., 2003; Miyao et al., 2001; Pillai et al., 2003; Rosenheck and Choder, 1998; Sheffer et al., 1999; Woychik and Young, 1989). To determine whether Rpb4 PTMs are biologically significant, we first determined whether they respond to temperature stress. Indeed, additional isoforms were detected in Rpb4 upon temperature shiftup (Fig. 2A), indicating that Rpb4 PTMs respond to changes in environmental conditions. The 2D profile of Rpb5 did not respond to the high temperature as clearly as Rpb4 (Fig. 2B); this profile helped us to orient the Rpb4 2D profiles of the two conditions.

**Fig. 2.**
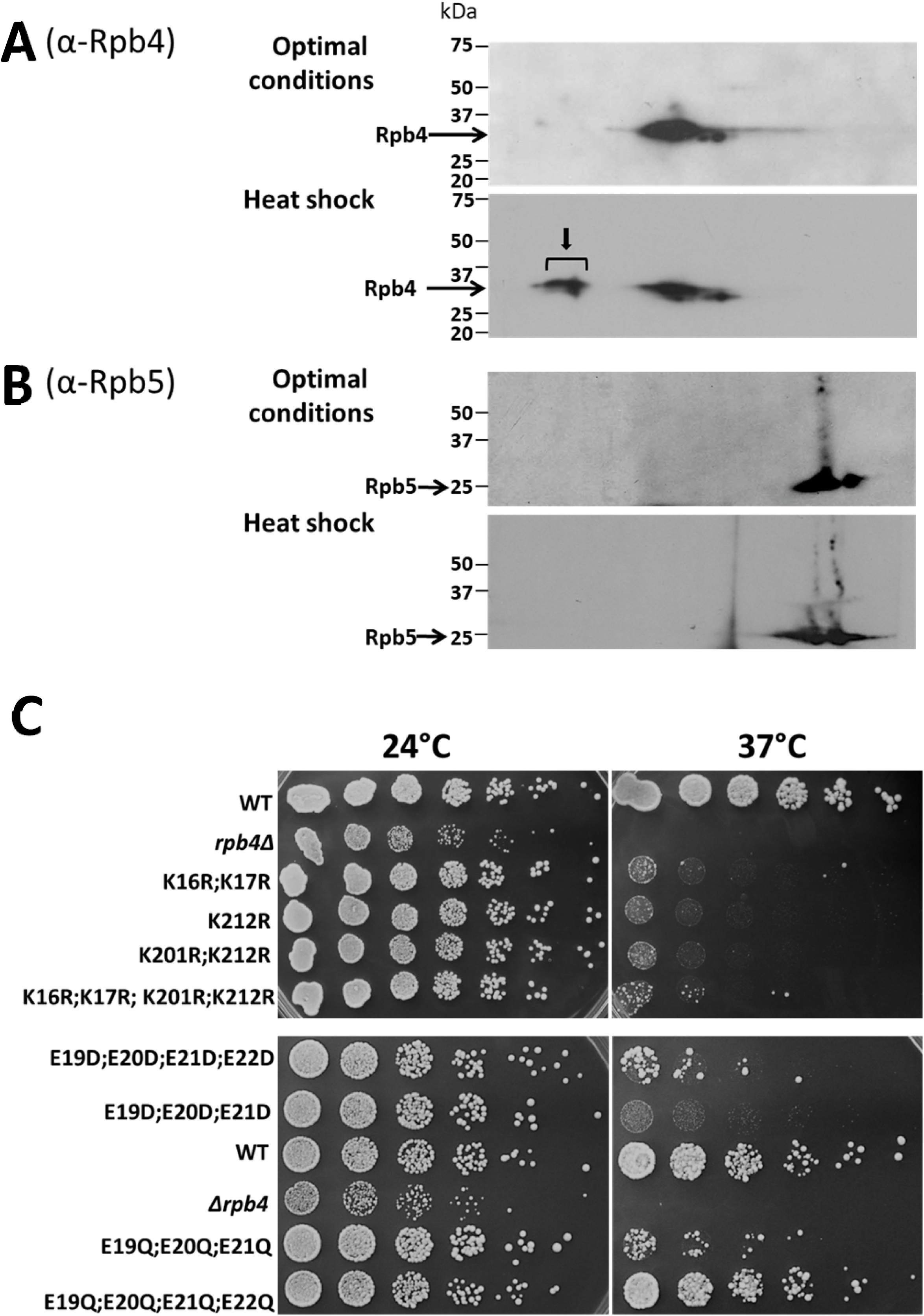
Mutations in Rpb4 residues that undergo PTMs compromise proliferation at 37°C – temperature that stimulates rapid change in Rpb4 isoform profile. (A-B) Rpb4 and Rpb5 isoform profiles obtained from WT strain (ySR007) expressing TAP-tagged *RPB4* under optimal conditions at 30°C or following a shift to 37°C. A culture of optimally proliferating cells was divided into two identical portions. One was harvested without heat shock and the other one was heat shocked by rapidly shifting to 37°C followed by shaking at 37°C for 20 minutes before harvest. Rpb4/7 was purified and equal amount of protein was analyzed as in Fig. 1C. The two membranes were reacted together with anti-Rpb4 antibodies and were expose to X-ray film together. Arrow points at stress specific spots. The membranes were then reacted, together, with anti-Rpb5 and exposed together to X-ray film. The Rpb4 spots remained faintly detectable (not detected in this short exposure). We then aligned the two films (the “optimal conditions” and “heat shock”), using Rpb5 as the reference point. This helped us to orient precisely the Rpb4 2D profiles of the two conditions. (C) Proliferation of WT and mutant cells at 24°C and 37°C. Optimally proliferating strains, indicated on the left, were diluted in a series of 5-fold dilutions and spotted on two YPD plates (first spot contained 3×10^4^ cells). Photos were taken after 3 days at the indicated temperature. See also Figs. S1, S3 and S4.

Next, we mutated a set of modified residues (Fig. S3) and found that the mutations affected cell proliferation at 37°C (Fig. 2C), indicating that the concerned modifiable residues are important for cell proliferation. Interestingly, mutating both Glutamic acid to Aspartic acid and Glutamic acid to Glutamine, which mimics methylation (Bornhorst and Falke, 2000; Endres et al., 2008), compromised proliferation at 37°C, suggesting that both states are important, most probably temporarily. Although, in principal, aspartic acid can become methylated, the temperature sensitivity of cells carrying E19-21D suggests that they cannot acquire the transient methylation required for efficient proliferation at 37°C. In contrast with 37°C, under otherwise optimal conditions at 24°C, all these mutations had little effect on cell proliferation. This is similar to the relatively little effect of *rpb4*Δ on cell proliferation at 24°C (Fig. 2C).

Collectively, the temperature sensitivity of the studied mutants, despite being highly conserved, and the response of the isoform profile to high temperature suggest that the different Rpb4 isoforms are biologically important.

### Rpb4 isoform profile is relatively robust, but is substantially affected by mimicking methylation of the E19-22 motif

To determine the relationship between the PTM residues, identified by MS, and the isoform profile, we mutated these modified residues. Fig. S4 shows that changes in steady-state Rpb4 protein levels never exceeded ±20% with respect to the WT. In previous work we down-regulated Rpb4 level by ~50% relative to the WT without affecting the cells’ capacity to proliferate at 30°C or 37°C (Duek et al., 2018). Given that here we observe smaller relative changes, we conclude that any possible effect on cellular functionality due to the mutations introduced to Rpb4 is not a consequence of their effects on Rpb4 protein levels.

We next performed 2D analyses of the mutants and discovered that the isoform profile is relatively robust to changes because most mutations did not affect the overall profile. Some conspicuous changes in distinct isoforms were observed (marked by arrows in Fig. 3A). For example, in the *rpb4*^K16R; K17R^ mutant, one isoform (a 2D spot) disappeared (marked by an arrow) and another isoform of a more basic protein appeared. In the *rpb4*^K212R^ mutant, a new spot appeared to the right, but also a new spot to the left. These relatively small changes demonstrate that the isoform profile is robust and most of the PTMs are not affected by these mutations except, perhaps, the isoforms that are specific to the mutated residues. However, interfering with the methylation capacity of the E19-22 motif compromised this robustness. Whereas replacing E with D had a small effect, replacing E with Q changed the profile substantially (Fig. 3B) as profiles became more dispersed with new spots appearing on the right and the left. Interestingly, despite the observation that replacing all four E with D had a relatively small effect on the isoform profile, we detected a severe effect of these mutations on cell proliferation at 37°C (Fig. 2C). This seeming discrepancy suggests that E19-22 methylation is transient yet plays an important role in Rpb4 functionality. Accordingly, replacement of all E19-22 with D could not be observed by 2D because it represents the ground state of the isoform profile, whereas the methylated state is transient and does not affect the visible profile; only upon replacing E with Q is the effect visible as it loses its transient nature. Moreover, the substantial change of the profile upon replacing E with Q suggests that the E19-22 motif regulates other Rpb4 PTMs. Collectively, the Rpb4 isoform profile is robust, but mutations that mimic methylation of the E19-22 motif seem to play a key role in shaping it. These results encouraged us to further investigate the function of this motif and the impact of its transient methylation on Rpb4 function.

**Fig. 3.**
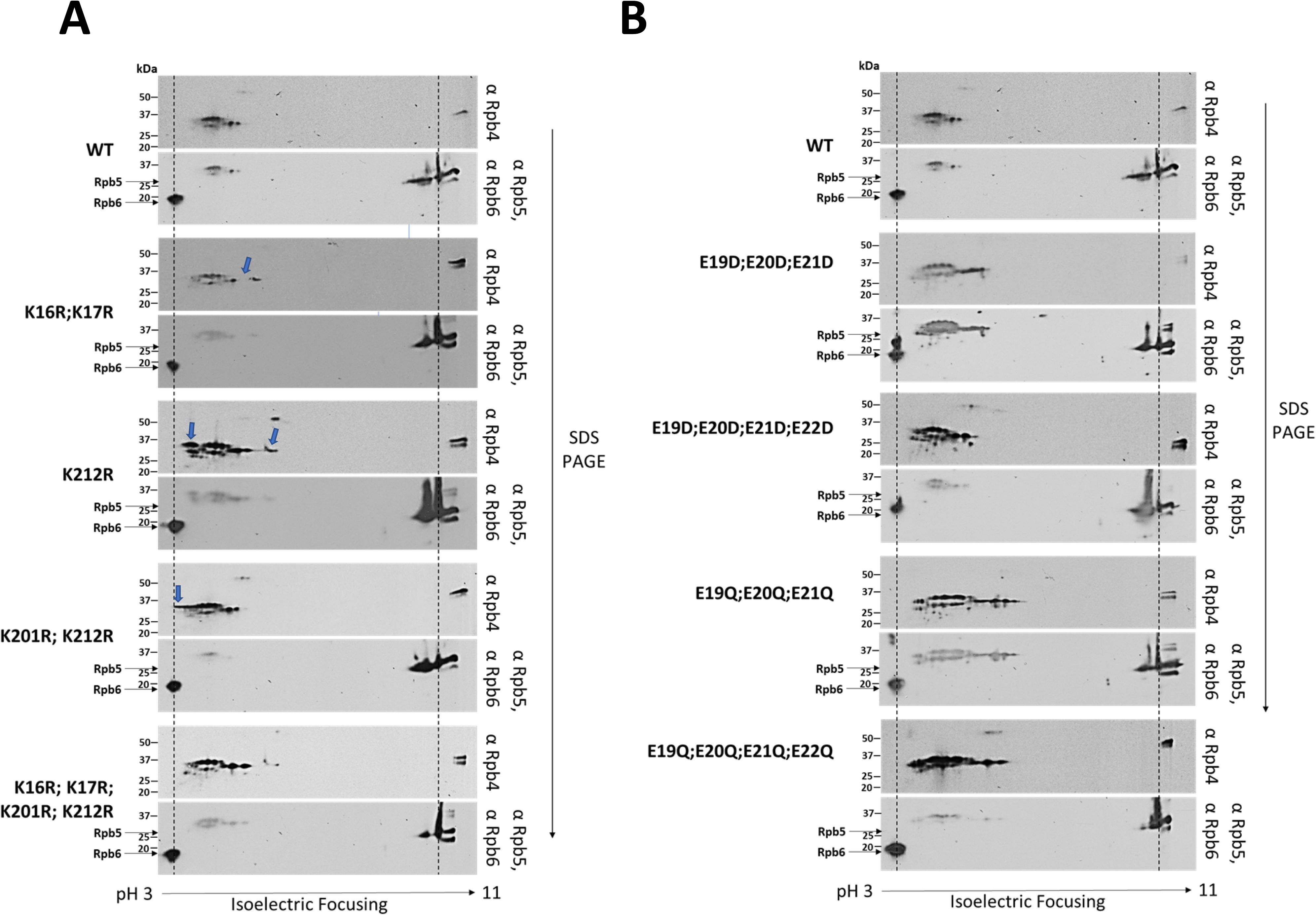
Rpb4 isoform profile is relatively robust but is shaped by methylation of the E19-22 motif. Rpb4 was tandemly affinity purified and equal amount of protein was subjected to 2D gel electrophoresis followed by western blot analysis as described in Fig. 1C. After probing the membranes with anti-Rpb4 Abs, they were probed with anti-Rpb5 and anti-Rpb6 Abs. These two proteins have opposing extreme pKa and therefore they demarcate the two ends of the membranes. The signal of the anti-Rpb4 (mouse Abs) is weekly detected after reacting the membrane with anti-Rpb5/6 (rabbit Abs). This helped us to unambiguously align Rpb4 spots. The dashed lines are shown to highlight spots that were used in the alignment. (A) Mutation in K16, K17, K201, K212. Arrows point at clear noticeable mutant-specific spots. (B) Mutations in the E19-22 motif.

### Rpb4 modifications are dependent on normal recruitment of Rpb4 to Pol II

We have previously shown that binding of Rpb4 to Pol II transcripts and its capacity to modulate mRNA translation and decay is dependent on its binding to Pol II (Goler-Baron et al., 2008; Harel-Sharvit et al., 2010). This provoked us to propose that co-transcriptional binding of mRNAs with some proteins including Rpb4/7 (“mRNA imprinting”) plays an important role in the cross talk between transcription and post-transcriptional stages (Choder, 2011). To determine whether Rpb4 modifications occur in the context of the mRNA/Rpb4 complex, we used an *rpb6^Q100R^* mutant strain, as its Pol II loses ~75% of its Rpb4/7 binding capacity (Tan et al., 2003). Rpb4 in this strain binds mRNAs poorly (Goler-Baron et al., 2008). The isoform profile of Rpb4 purified from *rpb6^Q100R^* mutant cells was substantially different than that of the WT. Whereas in WT cells 20 major spots of Rpb4p were detected (Fig 4, WT panel), only 7 were detected in the mutant (Fig. 4, *rpb6*^Q100R^ panel), among which spots # 4, 5, 6 and 7 increased 2–5 fold. Thus, except for 4 spots that exhibited increased intensity, all others were substantially decreased or disappeared. We propose that most modifications require Rpb4 binding to Pol II; moreover, as Rpb4 binding to mRNA occurs only in the context of Pol II, and it is compromised in *rpb6^Q100R^* cells (Goler-Baron et al., 2008; Harel-Sharvit et al., 2010), we suspect that most modifications occur in the context of mRNA/Rpb4/7. This possibility was supported by our observation that the nature of Rpb4 PTMs changes as the mRNA/Rpb4/7 complex progresses through its various stages (e.g., transcription, translation; see next section) and by the impact of Rpb4 PTMs on mRNA degradation (see below).

**Fig. 4.**
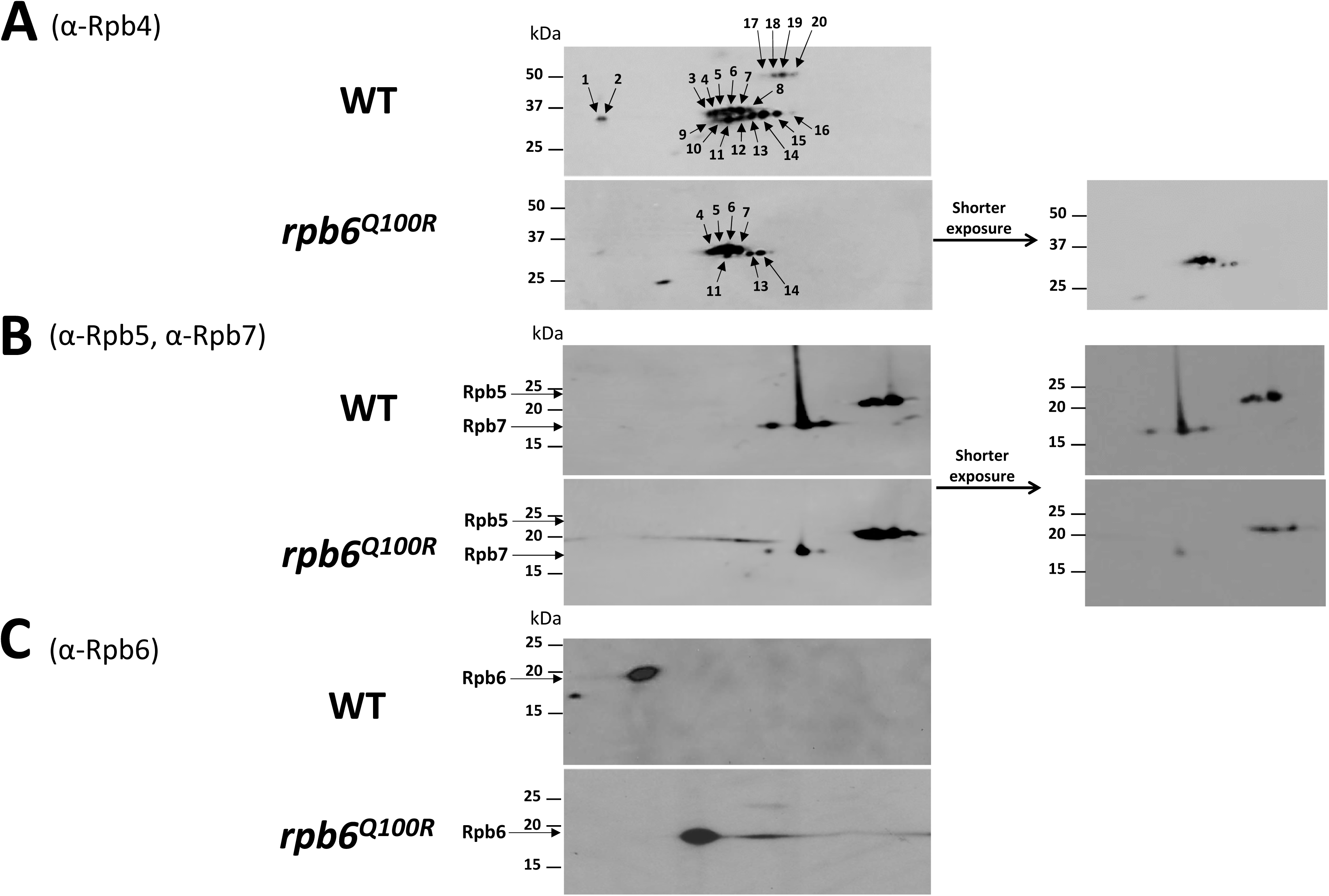
Isoform profile of Rpb4 is affected by Rpb4/7 recruitment to Pol II. Rpb4-TAP was tandemly affinity purified from extracts of WT (yVG54) or *rpb6*^Q100R^ (yVG55) cells. Although Rpb4/7 level in the mutant cells is comparable to WT, Rpb6^Q100R^-containing Pol II poorly recruits Rpb4/7 (Goler-Baron et al., 2008; Harel-Sharvit et al., 2010; Tan et al., 2003). Equal amount of purified protein was analyzed by 2D gel electrophoresis followed by western blotting analysis as in Fig. 1C. The two membranes were reacted together with either anti-Rpb4 Abs (A), or with anti-Rpb7 Abs and anti-Rpb5 Abs (B) or anti-Rpb6 Abs (C). The membranes were then exposed together to an X-ray film and to ImageQuant LAS 4010 Imaging & detection system (GE Healthcare Life sciences) for quantification. Rpb4 isoforms are marked, arbitrarily, with numbers. Quantification of spot intensities revealed that spots 4, 5, 6 and 7 increased 1.9, 4.7, 5.2, and 3.9 folds respectively, whereas spots 11, 14, 15 decreased 2.5-fold or more and the rest were undetectable. Note that due to replacement Q with R, Rpb6 isoelectric point changed. Rpb6 intensity demonstrated equal loading

### Different Rpb4/7 isoforms are temporarily associated with different complexes that control mRNA biogenesis, translation, and decay

Rpb4 functions in transcription (Babbarwal et al., 2014; Choder, 2004; Farago et al., 2003; Miyao et al., 2001; Pillai et al., 2003; Rosenheck and Choder, 1998; Runner et al., 2008; Verma-Gaur et al., 2008), mRNA export (Farago et al., 2003), Translation (Harel-Sharvit et al., 2010; Villanyi et al., 2014), mRNA degradation (Duek et al., 2018; Goler-Baron et al., 2008; Lotan et al., 2007, 2005; Schulz et al., 2014). This versatility is made possible by virtue of Rpb4 to physically interacting with key regulators of these stages of the mRNA lifecycle, among them are Pol II, the translation initiation factor 3 (eIF3), and the scaffold of the major cytoplasmic mRNA decay complex - Pat1 (Parker, 2012). We also reported that Rpb4 is involved in the cross talk between transcription and translation (Harel-Sharvit et al., 2010), and between transcription and mRNA degradation (Goler-Baron et al., 2008). To determine which of the isoforms are associated with each of these complexes, we affinity purified Pol II, eIF3, and Pat1 and determined which Rpb4 isoforms are associated with each individual complex. First, we tandemly purified the Pol II complex from the chromatin pellet, using TAP-tagged Rpb3. Specifically, cell extract was centrifuged, and two fractions were recovered: the chromatin pellet and the supernatant. The pellet was digested with DNase I, to release Pol II, and the digest was tandemly affinity purified (Gavin et al., 2002). Only 2 isoforms of Rpb4 and one isoform of Rpb7 could be co-purified with Rpb3-TAP (Fig: 5, “Pol II complex chromatin associated” panel). Interestingly, additional isoform of both Rpb4 and Rpb7 were co-purified (marked by arrows) with Rpb3-TAP from the supernatant fraction (Fig: 5, “Pol II complex not associated with chromatin” panel), suggesting that there are two distinct pools of Pol II, chromatin-bound and chromatin-free (not engaged in transcription) whose Rpb4/7 is differentially modified. Next, we purified eIF3 complex using Nip1, and detected three (relatively basic) distinct Rpb4 isoforms of and one Rpb7 isoform (Fig. 5, “Translation complex” panel). We subsequently purified the mRNA degradation complex using Pat1 and observed three isoforms of Rpb4 and one isoform of Rpb7, (Fig. 5, “Decay complex” panel). Two of these isoforms cannot be detected in purified Rpb4 alone (Fig. 5, “Total” panel), suggesting that these isoforms represent a minor population and are short lived. Taken together, each distinct stage of the mRNA lifecycle is associated with specific isoforms of Rpb4/7. These results indicate that several Rpb4/7 PTMs are transient and temporarily regulated.

**Figure 5.**
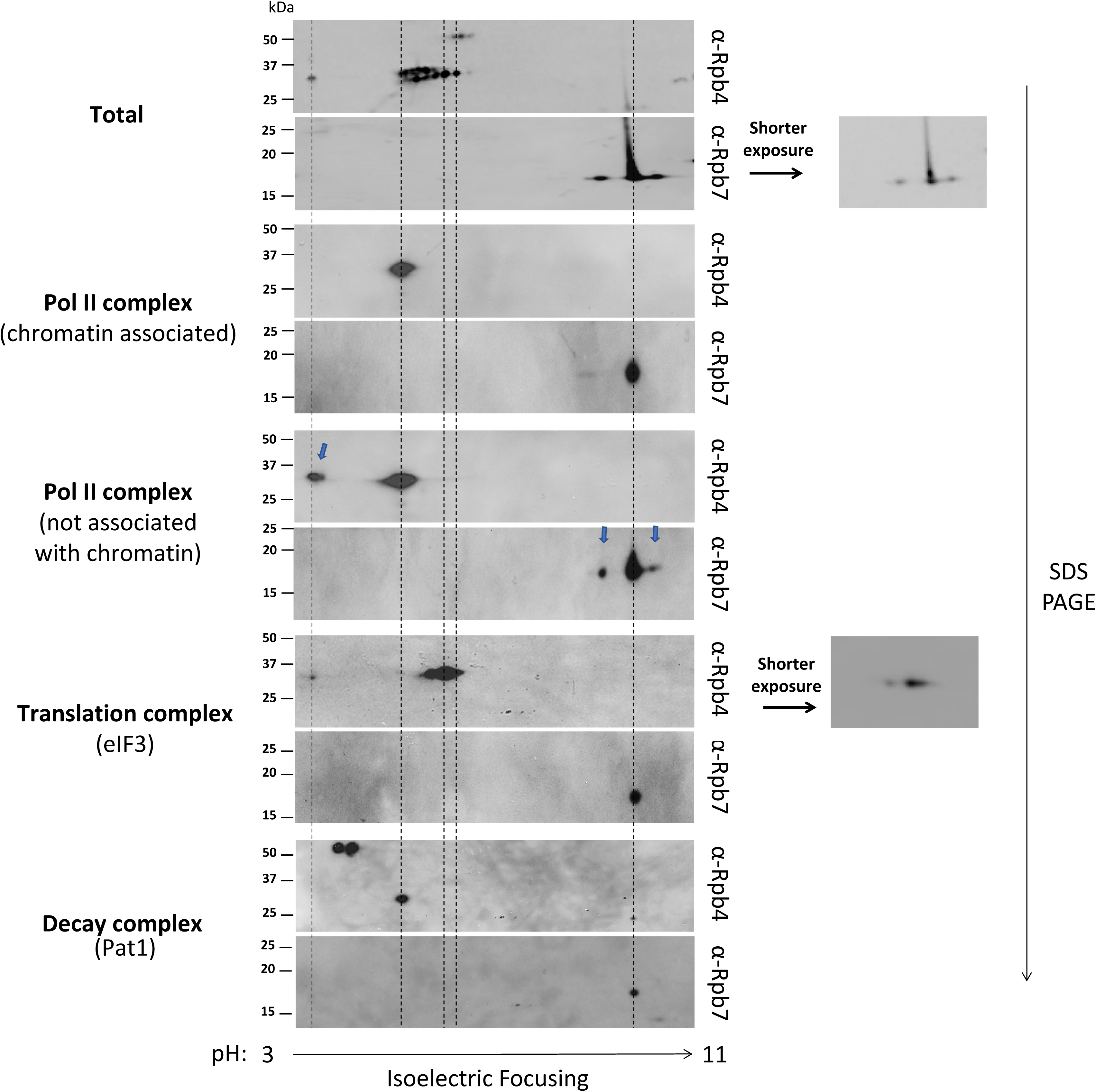
Different Rpb4 isoforms are associated with distinct stages of the mRNA life cycle. Isogenic strains expressing either Rpb4-TAP (“Total” panel), Rpb3-TAP (“Pol II complex” panels), Nip1-TAP (“Translation complex” panel) or Pat1-TAP (“Decay complex” panel) were grown in rich media, at 30°C and harvested at late log phase (~3×10^7^ cells/ml). Protein complexes were purified using the TAP procedure (Gavin et al., 2002). Chromatin associated or not associated complex were isolated as details in Methods. The proteins were subjected to 2D gel electrophoresis followed by western blotting analysis as in Fig. 1C. The membranes were reacted with anti-Rpb4 antibodies and later with anti-Rpb7, which was also used to align the Rpb4 spots profile, in the same manner as Rpb5 in Fig. 2B. The blue arrows point at spots that were specific to Rpb4 and Rpb7 that interacted with chromatin-free Pol II (probably not engaged in transcription).

### Mutations in Rpb4 E19-22 motif affect Rpb4’s interactome

By tandem affinity purification, we purified Rpb4 from WT cells and from cells carrying mutations in the E19-22 motif and analyzed them by mass spectrometry (MS). For identification of protein interactors, MS data from three biological replicates was analyzed using the MaxQuant program. Statistical analysis (mainly t-tests) was done using Perseus software and the results were expressed by volcano plots, as detailed in the caption of Fig. 6. In all our samples, save the control, we affinity purified comparable amount of Rpb4 peptides (Table S1). We note that since Rpb4 transiently interacts with several complexes, many of the interacting Rpb4 isoforms represent minor populations and therefore were not identified using the statistical analyses that we employed. For example, known interacting partners, including Pat1, Ccr4, Lsm2, Nip1 and Hcr1 (Babbarwal et al., 2014; Lotan et al., 2005; Miller et al., 2018) (Harel-Sharvit et al., 2010) were not identified as significant hits in our analysis. On the other hand, many known interactors of Rpb4, including all Pol II subunits, and transcription factors like Tfg1, Tfg2, Sua7, Taf14, Iwr1, the two subunits of the capping enzyme Cet1 and Ceg1, and Nab3 (Allepuz-Fuster et al., 2014; Chung et al., 2003; Czeko et al., 2011; Krogan et al., 2006; Mosley et al., 2013) were identified as significant hits in WT cells (Fig. 6A). These proteins serve as internal controls attesting to the efficient pull-down of Rpb4. Our data indicate that all Rpb4 mutant forms bind Pol II comparably to WT (Fig. S5, Table S1). Moreover, we found that all the studied mutant forms of Rpb4 bind Rpb7 comparably to WT (Fig. S5, Table S1). Importantly, then, the effects of the studied Rpb4 PTMs on cell proliferation (Fig. 2C), mRNA synthesis and decay (see next section) are not related to Rpb4-Rpb7 interaction and to the recruitment of Rpb4/7 to Pol II. We further have identified new Rpb4 interactors, such as the cap-binding protein Cbc2, a subunit of the exosome Rrp46, a component of the Ccr4-NOT complex Not1 (Cdc39) (although, interestingly, not other components of this complex), Rtt103, and the FACT subunit Spt16. We note that our approach could not differentiate between a direct and indirect interactions. Nevertheless, we can conclude that specific interactions affected by perturbing the E19-22 motif, discussed below, are not mediated by Pol II because all our mutant forms interact with Pol II comparably to WT (Fig. S5, Table S1). Interestingly, WT Rpb4, and in fact all the studied mutant forms, bind the largest subunit of Pol I Rpa190. Since no other Pol I subunits were co-purified, Rpb4 probably binds Rpa190 outside the context of Pol I.

**Fig. 6.**
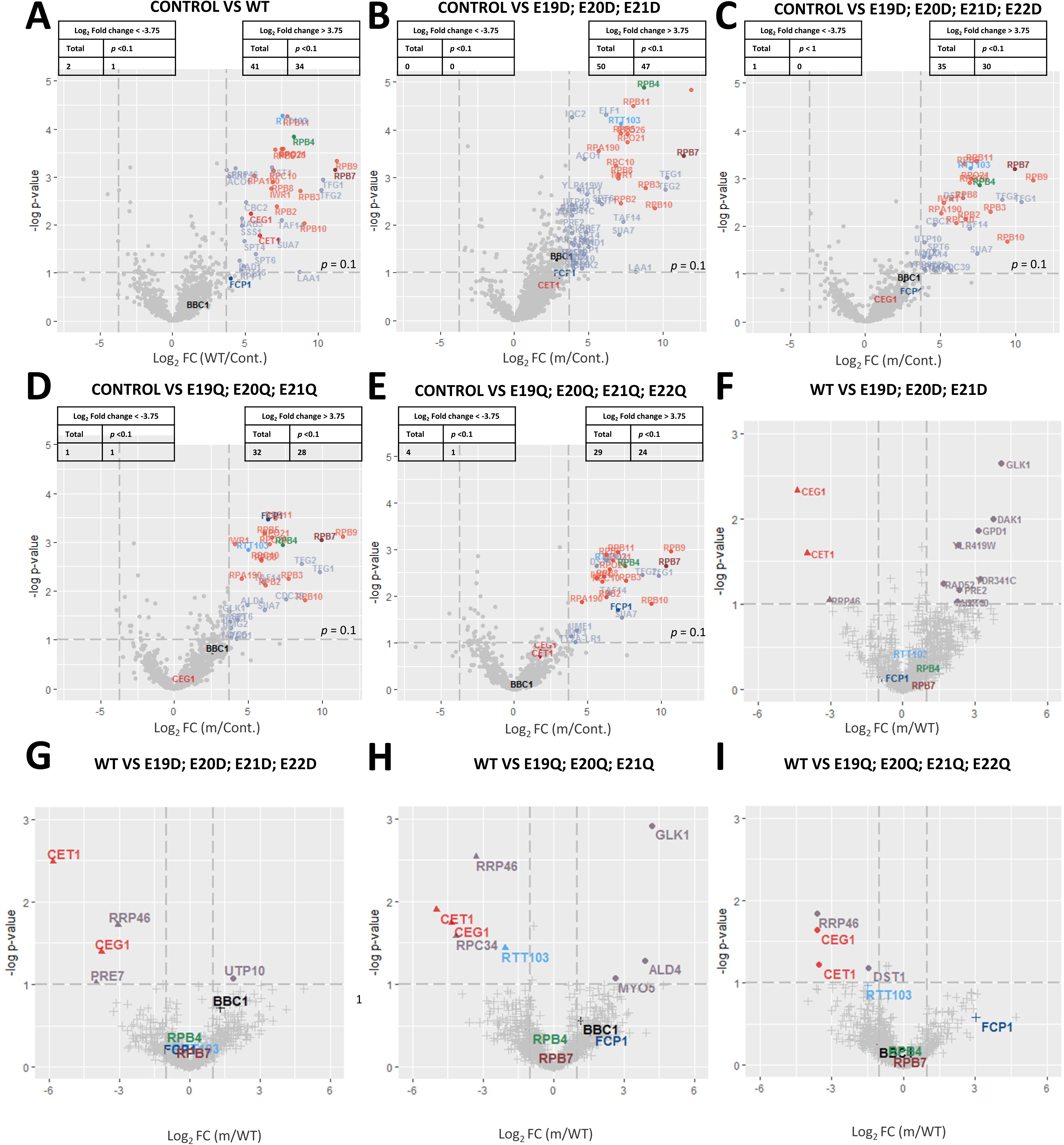
Mutations of Rpb4 E19-22 residues affect Rpb4’s Interactome. MS analyses of 3 biological replicates of cells carrying either no-tagged (control) or the indicated TAP-tagged Rpb4 were performed as detailed in Methods. The figure shows volcano plots comparing different groups, as indicated on top of each plot. Cont. - control; m - mutant (the mutations are indicated at the top of each plot). (A-E) Log_2_ fold changes (FC) in the levels of proteins co-purified with Rpb4-TAP (A) or its mutant forms (B-E), relative to those in the no-tagged control, was plotted relative to the significance. Pol II subunits are highlighted in red. (F-I) Log_2_ FC in the levels of proteins co-purified with the indicated mutant forms, relative to the Rpb4-TAP (WT), was plotted relative to the significance. Among the significant hits, only those that also were significantly enriched with respect to the no-tagged control are shown here. Specifically, in case that the WT protein was enriched over the mutant one, we used analyses shown in A; in case the mutant protein is enriched over the WT one, we used analyses shown in B-E. See also Figs. S4 and S5.

We then examined whether perturbing the Rpb4 E19-22 motif affected the Rpb4 interactome. Comparing the repertoire of proteins that were co-purified with WT Rpb4 and Rpb4 mutant forms has revealed a number of interesting observations. First, the interactions of both subunits of the mRNA capping enzyme and the exosome subunit Rrp46 are significantly compromised due to any perturbation of the motif (Fig. 6F-I and Table S1), suggesting that the interaction of Rpb4 with these proteins requires a certain combination of E19-22 methylation, which is missing in any of our mutants. Note that there are 2^4^ possible PTM combinations of the motif, most are not represented by our mutants. For example, consider a hypothetical possibility where efficient binding of these proteins with Rpb4 requires methylation of E19 while all other combinations of methylation disfavor interaction; such a specific combination is not present in any of our mutants. Regulated binding of the exosome subunit Rrp46 to Rpb4 suggests that the latter is capable of recruiting the exosome to either Pol II or Pol II transcripts and stimulating RNA degradation, consistent with a role we found for Rpb4/7 in 3’ to 5’ decay (Lotan et al., 2007). Second, Dst1 (TFIIS) interaction with Rpb4 is significantly compromised in E19-21Q and E19-22Q mutant cells (Fig. 6H-I and Table S1), suggesting that methylation of this motif releases Dst1 from Rpb4 binding. Dst1 stimulates RNA cleavage of backtracked Pol II (Nudler, 2012), thus playing a key role in regulating Pol II processivity and elongation efficiency. Indeed, deletions of both *DST1* and *RPB4* are synthetically lethal (Wery et al., 2004). The possible PTMs regulated interaction with Rpb4 suggests that Rpb4 stabilizes the interaction between Dst1 and Pol II in a reversible manner. Third, the interactions of Rpb4 with Rtt103 and Cbc2 are compromised due to mimicking constitutive methylated state in Rpb4 E19-21 (Fig. 6H). Note that all mutant forms bind Rtt103 (by comparing each mutant strain to the control lacking a tagged Rpb4), including the methylated mutants; however, comparison of WT with the mutant that mimics constitutive methylation at E19-21 revealed a significant difference (Fig. 6H-I and Table S1).

Mutants with glutamine in the E19-22 motif can be instrumental in identifying proteins that interact with a minor population of Rpb4 molecules that transiently carry methyl(s) in the motif. Our mutation strategy converted these minor populations into major ones and permitted the identification of these interacting partners. For example, identification of the type I myosin motor protein Myo5, involved in actin polymerization, required E19-22Q configuration (Fig. 6H). This raises the possibility that Rpb4 interacts with actin cables and that this interaction requires methylation of the motif. Consistently, *RPB4* interacts genetically with *ACT1*, encoding actin (Collins et al., 2007). We note that several other actin-related proteins were identified as potential interacting partners but did not pass our stringent cutoff.

Collectively, the E19-22 motif of Rpb4 influences Rpb4’s interactome. Importantly, the interactions of Rpb4 with some transcription-related factors are affected by mutations that mimic methylation. However, in the context of just Pol II, we identified only two major Rpb4 isoforms (Fig. 5, “Pol II complex”), suggesting that methylation of the E19-22 motif is transient and the specific isoforms responsible for the observed interactions escaped detection by the 2D profiling. A corollary of this conclusion is that the interaction of Rpb4 with these transcription related factors is also transient. Some interactions require PTMs that mimic methylation of E19-22, e.g., Myo5. Because we could not detect these factors among the significant hits of our WT strain, we propose that, also in these cases, both the specific PTMs and specific interactions are transient and represent a minor population among the Rpb4 molecules. It is worth noting that since modifications of the E19-22 motif affect the entire isoform profile (Fig. 3B), it is possible that the effects we detected are not direct but via the impact of the E19-22 motif on other modifications. However, because the motif protrudes outside of the Rpb4 structure and, in the context of Pol II, is facing the solvent (Fig. S2C-E), we nevertheless suspect that the motif directly participates in at least some of the interactions.

### Mutations of the Rpb4 E19-22 motif affect mRNA buffering

Rpb4 plays important roles in transcription (Allepuz-Fuster et al., 2014; Miyao et al., 2001; Orlicky et al., 2001; Pillai et al., 2003; Rosenheck and Choder, 1998; Schulz et al., 2014; Verma-Gaur et al., 2008) and in mRNA degradation (Goler-Baron et al., 2008; Lotan et al., 2005; Schulz et al., 2014). We therefore investigated whether the E19-22 motif plays roles in these two opposing activities, the balance of which determines steady-state mRNA levels. The levels of many mRNAs are buffered by a feedback mechanism that minimizes changes in mRNA levels, defects in which can lead to abnormal levels (Haimovich et al., 2013b; Singh et al., 2019; Timmers and Tora, 2018). To determine whether the E19-22 motif is involved in mRNA buffering, we examined mRNA levels in strains carrying mutations in this motif. Since these mutations affect cell proliferation at high temperature, but not at 24°C, and because the mutant cells cannot proliferate at 37°C (Fig. 2C), we extracted RNAs from cells that had been proliferating for many hours at a relatively high, but not lethal, temperature (33°C), and determined the steady state mRNA levels by RNA-seq; no *rpb4*Δ strain was included in this experiment because it cannot proliferate at this temperature (Choder and Young, 1993). A mixture of one-hundred poly(A)+ RNA spike-ins was added to 500 ng of total RNA before library preparation, to be used for normalization. Utilizing DESeq2, we compared mRNA levels between both mutants and WT samples as well as between mutants. As shown in the volcano plots (Fig. 7A), mutating the E19-22 motif resulted in highly significant (see p-values) increases in most mRNAs (Fig. S6A), while E19-22D had a weaker effect than E19-21Q (Fig. 7 cf A and B, and Fig S6A). This unexpected increase of mRNA levels contrasts with the decrease in mRNA levels that characterizes *rpb4*Δ cells (Lotan et al., 2005; Schulz et al., 2014). We note that the levels of these Rpb4 mutant forms were comparable to Rpb4 level in WT cells (Fig. S4) and that they were normally recruited to Pol II (Fig. S5). Thus, these results suggest that the E19-22 motif is involved in mRNA buffering.

**Fig. 7.**
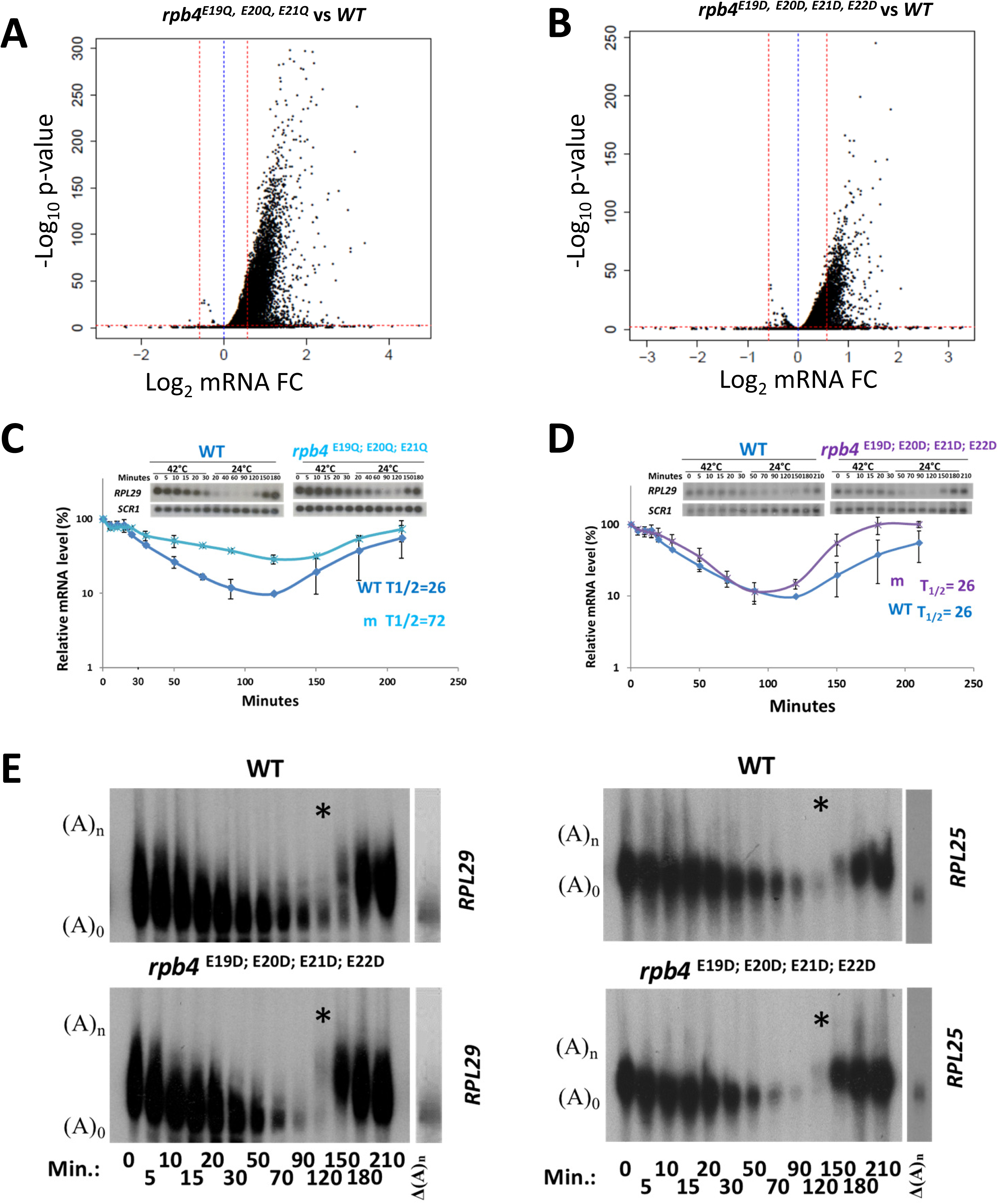
Mutations in Rpb4 E19-22 motif affect mRNA buffering. (A-B) Volcano plots showing fold changes of mRNA levels in the indicated mutant relative to those in WT cells, plotted relative to their significance [Log_10_(p-value)]. Three replicates of WT and the indicated mutant cells were cultures on rich medium (YPD) at 30°C till early mid-log phase (5×10^6^ cells/ml). Cells were shifted to 33°C and incubated for 6 h. RNA was extracted while cells were still at logarithmic phase (1.5 - 2×10^7^ cells/ml). Spike-in were added to 500 ng of RNA and RNA-seq was performed as detailed in Methods. Fold changes (FC) were calculated as the Log_2_ of the ratio of the expression value of each gene after normalization to the spike-in (~100 poly(A)+ RNAs) signals. (C-F) mRNA synthesis and decay kinetics. The indicated strains were cultured on rich medium (YPD) and allowed to proliferate at 24°C. Mid-log phase cultures were rapidly shifted to 42°C and incubated for 30 minutes at this temperature. The temperature shift blocked transcription (Lotan et al., 2007, 2005). The cultures were then rapidly shifted down to 24°C to permit transcriptional induction. Cell samples were taken just before the temperature shiftup (time 0) and at the indicated time points post temperature shiftup. (C-D) Standard Northern analyses. RNA, extracted from the indicated cell samples, was subjected to Northern analysis using the indicated radioactive probes, as described in Methods. Membranes were exposed to PhosphoImager screens and the amount of RNA was quantified using TotalLab Inc. software. The level of *SCR1* RNA, transcribed by Pol III, was used for normalization. Radioactive intensity, relative to that of *SCR1* mRNA, was normalized to time 0 that was defined arbitrarily as 100%. Error bars represents standard deviation from two biological replicates. For each biological replicate, three technical repeats were performed, thus a total of six repeats. Inset shows an example of one out of six repeats. The indicated mRNA half-lives (T_1/2_) were obtained from the slopes of the logarithmic curve. Note the logarithmic scale of the Y-axis. (E). PAGE Northern analysis. mRNA synthesis and decay analyses were done as in C-D except that the RNA samples were analyzed by the polyacrylamide Northern technique and probed with *RPL29* or *RPL25*, as indicated (see Methods). Lane “Δ(A)_n_” shows the position of fully deadenylated RNA, obtained by hybridizing mRNAs with oligo(dT) and digesting the hybrids with RNase H. Asterisk (*) marks the lanes showing the first time point where *de novo* transcription was observed in the mutant. A_0_ and A_n_, depicted on the left, denote the positions of deadenylated (A_0_) and poly(A) containing RNAs, respectively. Note that the newly transcribed transcripts (seen at time points 120-210 min.) run slower than the deadenylated RNAs (shown in the earlier time point) because of their long tail (Brown and Sachs, 1998), which helps identifying *de novo* transcription unambiguously. See also Fig. S6.

To determine which of the opposing mechanisms, mRNA synthesis or decay, was affected by E19-22 mutations, we shifted cells from 24°C to 42°C, which naturally blocks transcription of most genes (Lotan et al., 2007, 2005; Sheffer et al., 1999). After 30 min at 42°C, cells were shifted back to the low temperature. This stimulated transcription, which initiated after two hours of acclimation at this optimal temperature, allowing us to monitor transcription induction of these genes. Since most genes were affected, we chose genes that we had previously found to be affected by disrupting Rpb4 or Rpb7 (Lotan et al., 2007, 2005). Both mRNA synthesis and decay were adversely affected by replacing E with Q (Fig. 7C and S6B). In contrast, the E19-22D mutant degraded the studied mRNAs almost normally but synthesized them faster than WT (Fig. 7D and S6C).

Since abnormally fast transcription was unexpected, we examined transcription by other means, utilizing the different lengths of the poly(A) tails of decaying and newly synthesized RNAs. Specifically, the major pathway of mRNA decay starts with the shortening of the poly(A) tail. Thus, when we blocked transcription, RNAs gradually lost their tails and this could be observed by the polyacrylamide electrophoresis Northern technique (Decker and Parker, 1993; Lotan et al., 2007, 2005; Sachs and Davis, 1989). We chose to analyze *RPL29* and *RPL25* mRNAs because their short length enables detection of small differences in the sizes of their poly(A) tails. As shown in Fig. 7E, by 90 min. post-transcriptional arrest, *RPL29* and *RPL25* RNAs lost their tail. After this time point cells started to transcribe and to add long poly(A) tails to the nascent transcripts. Consequently, newly synthesized RNAs clearly migrated slower than the “old” deadenylated RNA that had existed before transcription initiated. We took advantage of this system to compare *de novo* transcription in WT and E19-22D cells. This approach clearly demonstrated that the mutant transcribed faster than WT (Fig. 7E). For example, at 120 min (lanes marked by *) the WT *RPL25* and *RPL29* RNAs migrated as deadenylated molecules, indicating that at this time point transcription was undetectable. In contrast with WT, in the mutant more than half of the RNA molecules migrated slower, due to a long poly(A) tail – a hallmark of *de novo* transcription. Enhanced transcription is illustrated also by the faster accumulation of poly(A)-containing *RPL29* and *RPL25* at later time points (150-210 min), as well as by the “standard” Northern analysis results (Fig. 7D).

Taken together, the abnormally high levels of mRNAs in the studied mutants indicate that the E19-22 motif is involved in buffering mRNA levels and that the underlying mechanism that broke the buffering function is affected by the type of mutations we introduced in the E19-22 motif. In particular, whereas E19-22D increases mainly transcription, the E19-21Q mutation affects both mRNA synthesis and degradation. Because our mutations differentially affected the two opposing functions of Rpb4 - mRNA synthesis and decay – the mutations uncoupled these functions. We conclude that, normally, Rpb4 executes two distinct functions in (i) mRNA synthesis and (ii) mRNA decay. Furthermore, the linkage between these functions is affected by mutations in its E19-22 motif. As we mutated the E19-22 motif in a manner that mimics its methylation, we propose that methylation of this motif is involved in the communication between mRNA synthesis and decay.

## Discussion

### Rpb4/7 carries numerous PTM combinations that change temporarily

Post-translational modifications play a part in coordinating the co-transcriptional assembly, remodeling and export of mRNPs (Tutucci and Stutz, 2011). Despite this significance, relatively little is known about their impact on the linkage between distinct stages of gene expression (e.g., mRNA synthesis and decay), or on cross talks between distinct machineries or functional complexes.

Here, we uncovered an unexpectedly high number of PTMs in a heterodimer (Rpb4/7) that functions in, and shuttles between, various machineries (see Introduction). Rpb4 exists in ~20 major isoforms and many more minor ones (Fig. 1D) while Rpb7 has 5 major isoforms; there are hence at least 100 possible combinations of Rpb4/7 isoforms. These PTMs are biologically significant because they are responsive to the environment, and mutations in the modified residues, even conserved ones (e.g., E to D, or K to R), affect cell capacity to respond to the environment. Remarkably, different repertoires of isoforms are associated with different machineries. More specifically, only two major isoforms of Rpb4 and one Rpb7 isoform were pulled down by chromatin-bound Pol II, whereas three different isoforms of Rpb4 were co-purified with eIF3 subunit Nip1 and three isoforms of Rpb4 and Rpb7 co-purified with Pat1 (Fig. 5). We have previously shown that Rpb4/7 binds Pol II transcripts co-transcriptionally, only in the context of Pol II, and then accompanies the mRNA throughout its life. Relevant to this study is our observation that compromising co-transcriptional mRNA binding, by using Rpb6^Q100R^, decreases Rpb4 binding with the translation complex as much as it compromises the function of Rpb4/7 in mRNA translation and decay (Goler-Baron et al., 2008; Harel-Sharvit et al., 2010) (Shalem et al., 2011). Using the same Rpb6^Q100R^ mutant form, here we found that most of the PTMs were dependent on prior recruitment of Rpb4 to Pol II (Fig. 4), and probably occur in the context of mRNA/Rpb4/7. Based on this and the different repertoires of isoforms that characterize distinct machineries, we propose that the mRNA/Rpb4/7 complex changes its PTMs temporarily as it progresses from one stage to the next. It seems that several isoforms found in the bulk Rpb4 were missing from these three complexes. For example, only one out of five Rpb7 isoforms were found bound to the three complexes that we examined. This suggests that the missing isoforms exist outside the context of Pol II, Nip1 and Pat1that we examined.

### E19-22 motif seems to be a hub that shapes the Rpb4 interactome

Several types of modifications have been identified by MS, as summarized in Fig. S1A. We mutated some of the residues that undergo modifications in Rpb4 and found that most compromised cell proliferation at 37°C (Fig. 2C), suggesting that these residues are critical for Rpb4 functionality. Interestingly, the Rpb4 isoform profile was found to be robust to most changes we made, except for local changes in specific isoforms, but mutations that mimic methylations of the E19-22 motif markedly reshaped the 2D profile. This motif, two E residues of which are highly conserved from yeast to human (Fig. S2B), resides in a loop that protrudes out of the heterodimer structure and seems to be accessible for interactions with outside factors (Fig. S2C-D). In the context of Pol II, the E19-22 motif is facing the solvent and can readily interact with outside factors (Fig. S2E). One possibility is that it is responsible for recruiting enzymes that modify other residues; however, we could not identify any modifying enzyme, suggesting either that these enzymes are not recruited by Rpb4 or that this recruitment is transient and escaped our detection. Unexpectedly, although replacing all four E with D had relatively small effect on the isoform profile, we detected a severe effect of these replacements on cell proliferation at 37°C (Fig. 2C). This seemingly discrepancy, combined with the substantial effect of the E to Q replacement on both the 2D profile and cell proliferation, suggests that E19-22 methylation is transient, and both the methylation and demethylation play important roles in Rpb4 functionality. We propose that replacement of E with Q, which mimics a constitutive methylation state (Bornhorst and Falke, 2000; Endres et al., 2008), disrupts the transient nature of the methylation state and compromises Rpb4/7 functionality.

Taken together, proliferation tests (Fig. 2C), the different repertoires of isoforms that characterize distinct machineries (Fig. 5) and MS results (Fig. 6) suggest that most PTMs are transient (for example, most of them are not found in the Pat1-associated Rpb4/7) and the combination of the transient methylation of theE19-22 motif is important for specific interactions and consequently for cell phenotype, and, as discussed below, for mRNA synthesis and decay.

### Mutations in the E19-22 motif affect mRNA buffering by uncoupling mRNA synthesis and decay

The role of Rpb4 in the linkage between mRNA synthesis and decay is controversial (see Introduction). Our observation that mutations in the E19-22 motif that mimic methylation affect both mRNA synthesis and decay are consistent with a model whereby any change in mRNA synthesis impacts changes in mRNA decay (Schulz et al., 2014; Sun et al., 2012). However, the increase in mRNA level indicates that the effect on transcription was not balanced by the effect on mRNA decay. Thus, mutations in Rpb4 break the mRNA buffering mechanism. Moreover, the E19-22D mutant affected only synthesis of the studied mRNAs but not their degradation (Fig. 7), indicating that Rpb4 plays two distinct functions that can be uncoupled by specific mutations. Being a Pol II subunit, one function is transcription. Some mutations in the E19-22 motif affect transcription, despite normal levels of these mutant forms (Fig. S4) and normal recruitment of Rpb4/7 to Pol II (Fig. S5). Thus, the main impact of these mutations seems to be related to the capacity of the motif to recruit certain transcription factors (see next section). The second function is mRNA decay, which is affected by replacements of the E19-22 glutamic acids with glutamine but not with aspartic acid. Since the mutations that compromise mRNA buffering mimic PTMs, transient PTMs of Rpb4/7 seem to play a role in the coupling between mRNA synthesis and decay.

### Possible new functions of Rpb4 in transcription-coupled RNA capping

Rpb4/7 is involved in all of the major transcriptional stages (see Introduction). Several hits from our MS approach are transcription factors known to interact with Rpb4/7, and some of these interactions were affected by the mutations we introduced in the E19-22 motif. Among them are two subunits of the capping enzyme (CE), the guanylyltransferase Cet1 and the triphosphatase Ceg1. Cet1 and Ceg1 interact with two Pol II domains, one of which is the phosphorylated Rpb1 C-terminal domain (CTD) and the other of which is located on the multihelical Foot domain of Rpb1 (Suh et al., 2010). More recent cryo-EM of a cocrystal of Pol II and recombinant Cet1 and Ceg1 revealed a small contact area between the two CE subunits and Rpb4/7, and cross-linking data further revealed a weak interaction between Rpb4 and Ceg1 (Martinez-Rucobo et al., 2015). It is not clear whether this interaction affects capping, either positively or negatively. Our results indicate that any of our mutations in the E19-22 motif compromised the interaction between Rpb4 and the two CE subunits. The simplest interpretation of these results is that a certain methylation pattern is required for efficient interaction, which is incompatible with any of our mutations. For example, if efficient binding of these proteins to Rpb4 requires methylation of just E19 while all other combinations of methylation disfavor interaction, this specific combination cannot be mimicked any of our mutants. The nature of the E19-22 mediated interaction of Rpb4 with the CE suggests that methylation of this motif is involved in regulating CE recruitment to Pol II, in addition to the regulation by Pol II CTD phosphorylation. Since the various mutations that similarly compromised Rpb4-CE interactions had different effects on transcription (Fig. 7 and S6), the defect in recruiting CE to Pol II cannot explain the defect in transcription that characterizes some of the mutants.

### Overview of the PTMs effects and a proposed model

Rpb4/7 protrudes away from the core of Pol II, thus potentially providing an independent landing pad for transcription related factors (Choder, 2004). Our results suggest that, indeed, Rpb4 serves as an additional landing pad for a number of proteins, some of which are known to also interact with the CTD (e.g., Rtt103). Moreover, the E19-22 motif is placed in a loop region that protrudes away from the bulk of the Rpb4/7 structure (Fig. S2C-E), providing a microenvironment accessible for interactions, in a manner regulated by temporal methylation. Despite our indications for numerous Rpb4 PTMS, in the context of just Pol II, we identified only two isoforms of Rpb4 (Fig. 5, Pol II complex). Thus, in the context of Pol II, the PTMs that regulate the Rpb4 interactome (e.g., those that affect its interactions with Rrp46, Cet1, Ceg2, Rtt103, Dst1) represent minor populations that are not detected by the 2D assay, most likely because they are transient. Consistent with transient interaction, the statistical analysis of our MS data did not identify Pat1, Lsm2, Ccr4, Hcr1 or Nip1 as significant hits, although previous works (Babbarwal et al., 2014; Harel-Sharvit et al., 2010; Lotan et al., 2007, 2005) and this work (Fig. 5, Nip1, and Pat1) clearly demonstrated that specific Rpb4 isoforms, which represent minor population of Rpb4 molecules (, interact with these proteins.

To overcome the transient nature of some of these interactions, we replaced some residues of the E19-22 motif with Q, thus mimicking constitutive methylation (Bornhorst and Falke, 2000; Endres et al., 2008). This approach, combined with MS, offered an opportunity to identify partners that potentially interact with the methylated E19-22 motif. An interesting hit was Myo5 (Fig. 6H, Table S1), a type I myosin motor. This unexpected result, which was not obtained with WT Rpb4, raised the possibility that regulated methylation of the E19-22 motif mediates mRNP transport along actin fibers.

An important feature of Rpb4/7 is its capacity to interact with distinct complexes that reside in different cellular localization such as Pol II, components of the cytoplasmic Pat1/Lsm1-7 complex and components of eIF3 (Choder, 2004; Harel-Sharvit et al., 2010; Kamenski et al., 2004; Lotan et al., 2007, 2005; Runner et al., 2008). We showed that Rpb4 needs to be recruited to Pol II in order to bind Pol II transcripts co-transcriptionally; Rpb4 then escorts these transcripts throughout their life while mediating major post-transcriptional stages (Duek et al., 2018; Goler-Baron et al., 2008; Harel-Sharvit et al., 2010). Here we show that the PTMs are also highly dependent on prior binding of Rpb4 and Rpb7 with Pol II (Fig. 4). Since we identified different Rpb4/7 isoforms in the context of each of these complexes (Fig. 5), we propose that some Rpb4/7 PTMs mediate the ability of Rpb4/7 to switch between these complexes and to properly function within any of the distinct complexes. As a proof of principle, we mutated the heavily modified E19-22 motif and found that mutations that mimic methylation adversely affect not only transcription but also mRNA decay.

A corollary of the new Rpb4 interactome is that the E19-22 motif is involved in binding several proteins. Steric considerations make simultaneous binding improbable. Our results, therefore, suggest that transient methylations of the E19-22 motif are involved in switching between the interacting partners. Binding of Rpb4 with Cet1/Ceg1, Dst1, Rtt103, exosome (Rrp46) and Myo5 are examples of this possibility. We propose that this small motif interacts temporarily and spatially with these proteins, thereby modulating (in due time) RNA capping, backtracking, transcription termination, exosome mediated decay and mRNA/Rpb4/7 transport. Previously, we proposed that Rpb4/7 functions as an mRNA coordinator, by integrating all stages of the mRNA lifecycle (Harel-Sharvit et al., 2010). Future work will be required to determine whether Rpb4/7 PTMs help coordinate not only mRNA synthesis and decay but, more globally, all stages of the mRNA lifecycle, thus integrating them into a system.

## Methods

### Yeast strains, growth conditions and construction of *rpb4* mutant strains

Yeast strains are listed in Table S3. Cells were grown in either YPD medium (2% Bacto Peptone, 1% yeast extract [Difco Laboratories], 2% dextrose), or synthetic complete (SC) medium lacking the appropriate amino acid as required. For all experiments, the inoculum was taken from cell cultures that were grown in log phase for at least seven generations. Mutations in *RPB4* were surgically introduced into the genomic locus *RPB4*. To this end, *RPB4* was first replaced with *CaURA3*, the mutant *rpb4* was then introduced by replacing *CaURA3*, utilizing homologous recombination and selection on 5-FOA. All mutants were verified by PCR followed by Sanger sequencing.

### Tandem Affinity Purification of protein complexes

Cell extracts were prepared from 4 liters of a yeast culture grown in YPD to late log phase (3−4 × 10^7^ cells/ml). Vacuum driven harvesting apparatus was used to harvest the cells. The harvested cells were immediately frozen in liquid nitrogen and stored at −80°C. Frozen cells were cryogenically ground (using liquid nitrogen) into powder by a mixer mill (Retsch MM 400) and stored at −80°C. Grindate was suspended in 13 ml of NP40 buffer (6 mM Na_2_HPO_4_, 4 mM NaH_2_PO_4_*H_2_0, 1% NP-40, 100 mM Potassium Acetate, 2 mM Magnesium Acetate, 50 mM Sodium Fluoride, 0.1 mM Na_3_VO_4_, 10 mM β-mercapto ethanol) containing protease inhibitors (1:25 Protease Inhibitor Cocktail [Roche], 2 mM PMSF, 50 µg/ml TLC-K, 2.5 mM Benzamidine, 2 mg/ml NaF, 1:100 Phosphatase Inhibitor Cocktail 2 [Roche], 1:100 Phosphatase Inhibitor Cocktail 3[Roche]) till the suspension was viscous, followed by centrifugation at 4°C for 20 minutes at 13000 rpm (21K g). In case of purifying the “Chromatin bound proteins” (Fig. 5), the pellet was resuspended with NP40 buffer till the suspension was viscous and treated with 2000 units of DNase I (Sigma # D7291). Equal amount of protein (normally 600 mg) was reacted IgG-Sepharose beads (GE healthcare; 350 μl of packed beads) and rotated for 2 hours at 4°C. The beads were transferred to a column and washed 3 times with 10 ml IPP150 buffer (10 mM Tris HCl pH=8.0, 100 mM Potassium Acetate, 0.1% NP40, 2 mM Magnesium Acetate, 10 mM β-mercapto ethanol, 1 mM PMSF), followed by 1 wash with TEV buffer (10 Mm Tris HCl pH=8.0, 100 mM Potassium Acetate, 0.1% NP40, 1 mM Magnesium Acetate, 1 mM Dithiothreitol). Cleavage was performed by adding 0.6 ml TEV buffer containing 100U TEV protease (self-made) and rotating at room temperature for 2 hours. The eluate (0.6 ml) was collected in a tube, followed by second elution with 1 ml of TEV buffer. Three volumes of Calmodulin Binding Buffer (CBB) buffer (0.1% NP40, 10 mM Tris HCl pH=8.0, 100 mM Potassium Acetate, 1 mM Magnesium Acetate, 1mM Imidazole, 2 mM CaCl_2_, 10 mM β-mercapto ethanol, 0.5 mM PMSF). CaCl_2_ was added to 1mM and a second purification was carried out overnight at 4°C using 300 μl of packed Calmodulin affinity resin (Agilent Technologies). The beads were transferred to a column and washed twice with 10 ml CBB buffer containing 0.1% NP40, followed by 1 wash with 10 ml CBB buffer containing 0.02% NP40. About 1 ml Calmodulin Elution Buffer (CEB) buffer (50 mM EGTA, 10 mM Tris HCl pH=8.0, 100 mM Potassium Acetate, 1 mM Magnesium Acetate, 0.02% NP-40, 1mM Imidazole, 10 mM β-mercapto ethanol) was added to the closed column. The closed column was incubated at 37°C for about 15 minutes before elution. Elutes were stored at −80°C.

### Two-dimensional Gel Electrophoresis

Protein sample was buffer-replaced to a sample solubilization buffer [7M urea, 2M thiourea, 2% w/v CHAPS, 65 mM DTT and 0.5% v/v ampholyte 3-10, and 3 µl Bromo-Phenol-Blue (BPB)]. Protein amount was quantified using Bradford analysis. The IPG strip (BioRad 3-10 NL Ready-Strip, 11 cm) was rehydrated with 200µg protein in 200 µl of the sample buffer. The rehydration was performed for 1 hour with no voltage and then for 10 hours with 50 mV. Isoelectric focusing was then performed for 1 h at 200 V, 1 h at 500 V, 1 h at 1000 V, 30 min linear increase to 8000 V and up to 12 h at 8000 V. Following Isoelectric focusing, each strip was equilibrated in 50 mg DTT in 5 ml of equilibration buffer containing 6 M Urea, 30% w/v glycerol and 2% SDS in 0.05 M Tris-HCl buffer (1.5 M Tris-HCl and 0.4% w/v SDS), pH 8.8 with 10 µl BPB for 15 min. A second equilibration step was performed for 15 min with 200 mg IAA in 5 ml of the above equilibration buffer. After equilibration, the immobilized pH gradient strips were loaded onto BioRad precast gels 4-20%, 11 cm and the proteins were separated along Bio-Rad’s dual color molecular weight markers for 2h at 100 V. Following the SDS-PAGE, proteins were transferred to a PVDF membrane (Hybond^P^; Amersham Biosciences). In some cases, following SDS-PAGE, the gels were stained with Coomassie.

### Western blot analysis

The membrane was blocked with fat-free milk for 1 hour at room temperature or over-night at 4°C. It was then reacted with primary antibody for 1 hour at room temperature, washed three times, each for 10 min, in TBS-T (20 mM Tris-Hcl pH=7.6, 150 mM Sodium Chloride, 1% Tween-80) and subsequently reacted with secondary antibody conjugated to Horse Radish Peroxidase (HRP). The membrane was washed three times in TBS-T. HRP activity on the membrane was detected with the Western Lightning Plus-ECL (Perkin Elmer) according to the manufacturer’s instructions.

### Identification of Rpb4 protein interactors and post translational modifications of Rpb4 by Mass Spectrometry (MS)

Affinity-purified Rpb4 samples were digested by trypsin, analyzed by LC-MS/MS on Q-Exactive plus (Thermo) and identified by Discoverer software against the *S.cerevisiae* proteome from the UniProt database, or against the Rpb4 protein sequence, and against a decoy database (to determine the false discovery rate). Sequest (Thermo) and Mascot (Matrix science) were the two search algorithms used. All the identified peptides were filtered with high/medium confidence, top rank, mass accuracy, and a minimum of 2 peptides. High confidence peptides have passed the 1% FDR threshold. Semi quantitation was done by calculating the peak area of each peptide. The area of the protein is the average of the three most intense peptides from each protein. For identification of protein interactors, MS data was analyzed using maxQuant program. Statistical analysis was done using Perseus software with the volcano plot option as follows: Label-free quant (LFQ) values of 3 MS analyses combining test strain and untagged *RPB4* control were transformed to log2 scale. After filtering out proteins that did not appear in 3 repeats of at least one of the groups (*RPB4* tagged or untagged control), missing values were imputed from normal distribution. The 2 groups were compared using t-test while applying permutation-FDR correction with FDR=0.05 and S0=0.1. In the volcano plot, x-axis represent the difference between the compared groups and the y axis represent the -Log(p value). For identification of post translational modifications, the identified high confidence peptides were screened for mass additions that represent acetylation (42.01057 Da), methylation (14.01565 Da), phosphorylation (79.96633 Da), and ubiquitination or neddylation (114.04293 Da). During sample preparation and subsequent manipulations, methanol was excluded to avoid *in vitro* methyl esterification.

### mRNA synthesis and degradation rate assay

The assay was performed as reported previously (Lotan et al., 2007, 2005). Briefly, transcription was blocked naturally by shifting cells that had been proliferating at 24°C for >7 generations to 42°C (when cell density reached 1× 10^7^ cells/ml). Cell aliquots were taken at the indicated time points, quickly chilled to ~4°C using liquid nitrogen (during 18 seconds), collected by 2 min centrifugation and flash-frozen in liquid nitrogen and stored at −80°C. After 30 minutes, the temperature of the culture was changed to 24°C by shifting the culture to the ice water and shaking vigorously for 48 seconds and then transferred to the 24°C incubator shaker. Cell aliquots were harvested as indicated in the figure legends, their RNA was extracted by hot phenol and equal amount of RNA was analyzed either by Northern blot hybridization or by PAGE Northern as described previously (Lotan et al., 2007, 2005).

### Determining mRNA level by RNA-seq

Cells were proliferated in three replicates in rich medium (YPD) for many generations till 5×10^6^ cells/ml and shifted to 33°C for 6 h. Cell were harvested at 1.5 – 2×10^7^ cells/ml. RNA was extracted by hot phenol technique. Quality control for total RNA was performed using TapeStation (Agilent). The RINe value of all samples was in the range of 9.6-9.8, indicating a high quality. 500 ng of total RNA was spiked in by adding 1ul of ERCC RNA Spike-In Mix (100 poly(A)+ RNAs) diluted 1:100 as specified in the protocol (Ambion, Thermo Fisher Scientific, cat no. 4456740). Nine RNAseq libraries were produced according to manufacturer’s protocol (NEBNext UltraII Directional RNA Library Prep Kit for Illumina, cat no. E7760) using 500 ng total RNA + the spike in. mRNA pull-up was performed using the Magnetic Isolation Module (NEB, cat no. E7490). All libraries were mixed into a single tube with equal molarity. The RNAseq data was generated on Illumina NextSeq500, 150 cycles (75 paired-end), mid-output mode (Illumina, cat no. 20024904). About 20 million mapped reads were obtained per sample. Quality control was assessed using Fastqc (v0.11.5), reads were trimmed for adapters, low quality 3’ and minimum length of 20 using CUTADAPT (v1.12). 75 bp paired-end reads were aligned to a yeast reference genome with ERCC spike-in sequences (Saccharomyces_cerevisiae.R64-1-1, ERCC http://tools.invitrogen.com/downloads/ERCC92.fa) and annotation file (Saccharomyces_cerevisiae.R64-1-1.96.gtf,https://github.com/NCBI-Hackathons/HASSL_Homogeneous Analysis of SRA rnaSequencing Libraries/blob/master/Spike-Ins/ERCC92/ERCC92.gtf) using STAR aligner (v2.6.0a). Differential expression analysis was performed using DESeq2. To normalize across samples, we first computed normalization factors for samples within the same condition via DESeq2’s ‘estimateSizeFactorsForMatrix’ function using all genes. This yielded intra-condition normalization factors (one per sample). Note that these factors only depend on samples from the same condition and thus purely account for variation among replicates. The intra-condition normalization factors were then applied to the spike-ins associated with each sample and the average value of each spike-in was taken within each biological condition. Normalization factors across conditions were then estimated as before but restricted only to averaged spike-ins, yielding a single inter-condition normalization factor for each condition. This allowed us to make use of the clear and consistent shifts in spike-in count levels across different conditions while being robust to intra-condition variation among spike-ins, preserving power. Finally, we obtained sample-specific normalization factors by taking the product of the appropriate intra- and inter-condition normalization factors and dividing by the geometric mean of these values. We then provided these as the SizeFactors for DESeq2’s differential expression algorithm.

## Acknowledgments

We would like to thank Gal Haimovich for critically reading the manuscript, the Genetic Center, Faculty of Medicine, Technion for preparing libraries and sequencing, A Sentenac for anti-Rpb7 Abs and the Choder’s group for stimulating discussions. This work was supported by the Israel Science Foundation (1472/15) to MC.

## Author Contributions

SR performed most of the experiments and wrote the manuscript. LG initiated the study and performed some of the 2D analyses. JF performed the statistical and computational analyses of the RNA-seq data, and edit the original draft of the manuscript. KB and TZ obtained the MS data. SU analyzed the MS data. MC conceived the study and wrote the manuscript.

## Supplementary figure legends

**Supplemental Fig. 1. Related to Fig. 2.**
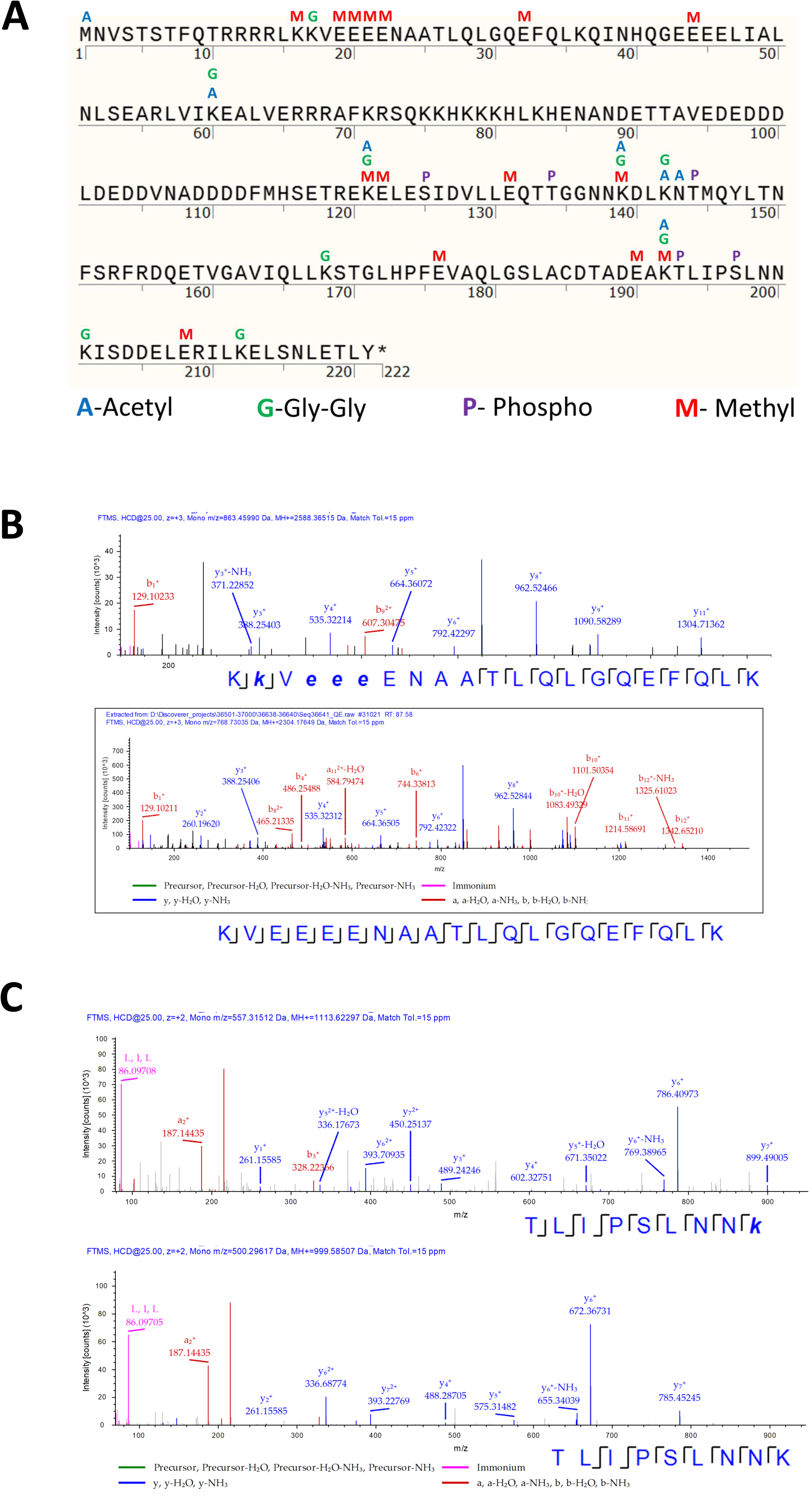
Mass Spectrometry analysis of Rpb4 reveals Post Translational Modifications. **(A)** Rpb4 sequence with identified modifications, indicated on top of the modified residue. Three biological replicates of affinity purified Rpb4 samples were digested by trypsin, analyzed by LC-MS/MS on Q-Exactive plus (Thermo) and identified by Discoverer software against the *S. cerevisiae* proteome from the UniProt database, against the Rpb4 protein sequence, and against a decoy database (in order to determine the false discovery rate). Sequest (Thermo) and Mascot (Matrix science) were the two search algorithms used. All the identified peptides were filtered with high/medium confidence, top rank, mass accuracy, and a minimum of 2 peptides. High confidence peptides have passed the 1% FDR threshold. (*FDR =false discovery rate, is the estimated fraction of false positives in a list of peptides). For identification of post translational modifications, the identified high confidence peptides were screened for mass additions that represent acetylation (42.01057 Da), methylation (14.01565 Da), phosphorylation (79.96633 Da), and ubiquitination or neddylation (114.04293 Da). (B) Annotated MS/MS spectra of a modified peptide containing the E19-22 Motif. The spectra display identified ions corresponding to the main fragmentation series obtained by collision-induced fragmentation of the peptide backbones and identified using the Sequest search engine. (C) Modified peptide with Gly-Gly addition on K2 and three methylations on E4, E5 and E6. Bottom panels of B and C show parallel unmodified peptide. (*y* ions are fragmentation products from the c-terminus and *b* ions - from the N-terminus).

**Supplemental Fig. 2.**
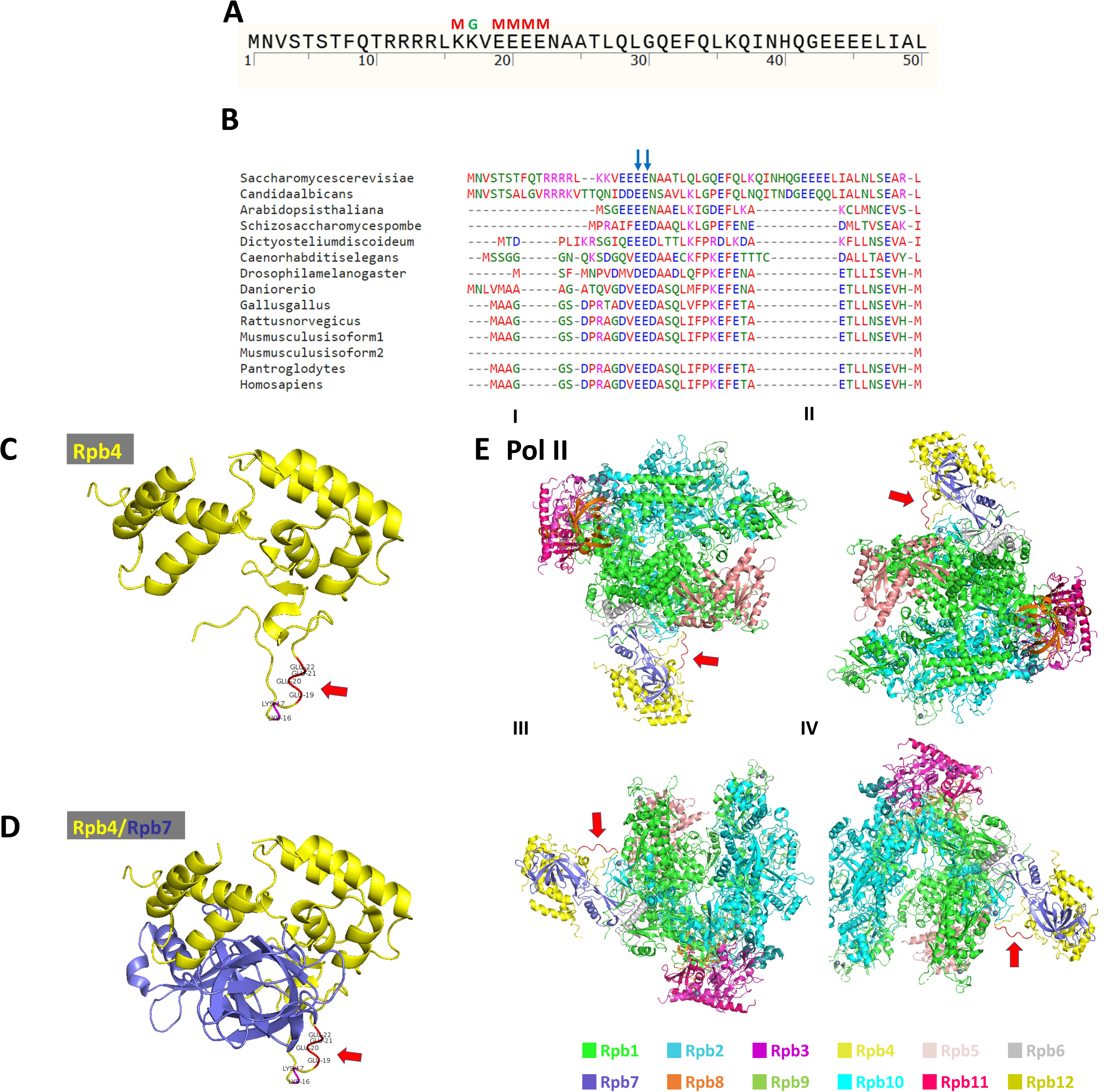
The Rpb4 E19-22 motif is conserved and located in a loop that protrudes out of Rpb4/7 structure, whereas in the context of Pol II structure it is facing the solvent. (A) Fifty residues of Rpb4 N-terminal. The E19-22 motif is indicated. (B) Multiple sequence alignment of the Rpb4 protein showing the highly conserved E21 and E22 residues, pointed by arrows. (C-D) The crystal structure of Rpb4 (C), or Rpb4/7 (D) (PDB ID 1WCM), in which the modified residues K16 to E22 are labelled in red and pointed by a red arrow. (E) Different angles and point of view (I to IV) of Pol II showing the protrusion of the Rpb4 loop (K16 to E22) that is facing the solvent. The modified residues are labeled as in C. All the structures were visualized using pyMOL program version 2.0.4.

**Supplemental Fig. 3. Related to Fig. 2.**
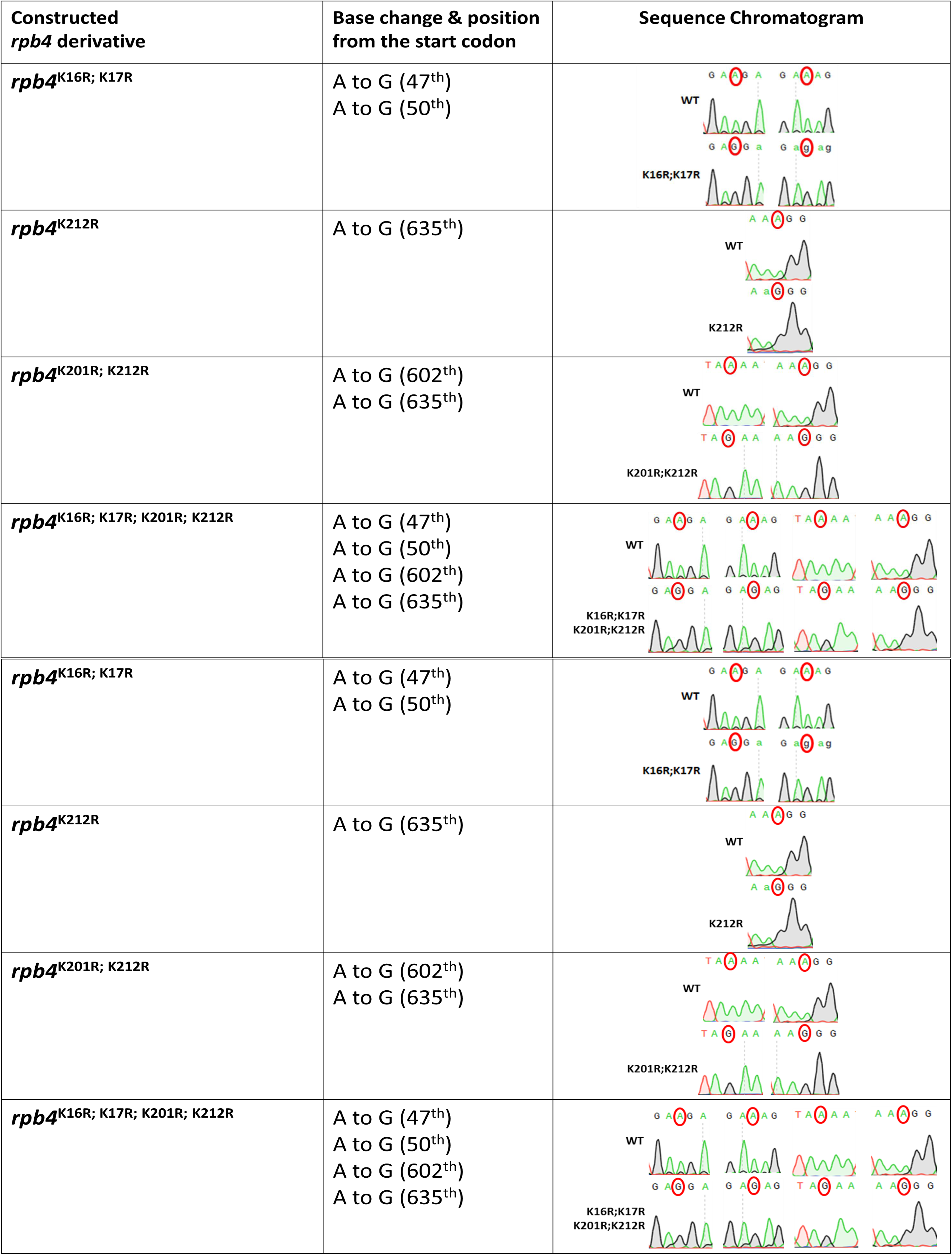
Introduction and verification of *RPB4* mutations. The table shows the mutated Rpb4 residues, the base pair changes involved and the verification of mutations by Sanger sequencing. The mutations were surgically introduced into the genomic *RPB4*. To this end, *RPB4* was first replaced with *CaURA3*, the mutant *rpb4* was then introduced by replacing *CaURA3*, utilizing homologous recombination and selection on 5-FOA.

**Supplemental Fig. 4. Related to Fig. 2 and 6.**
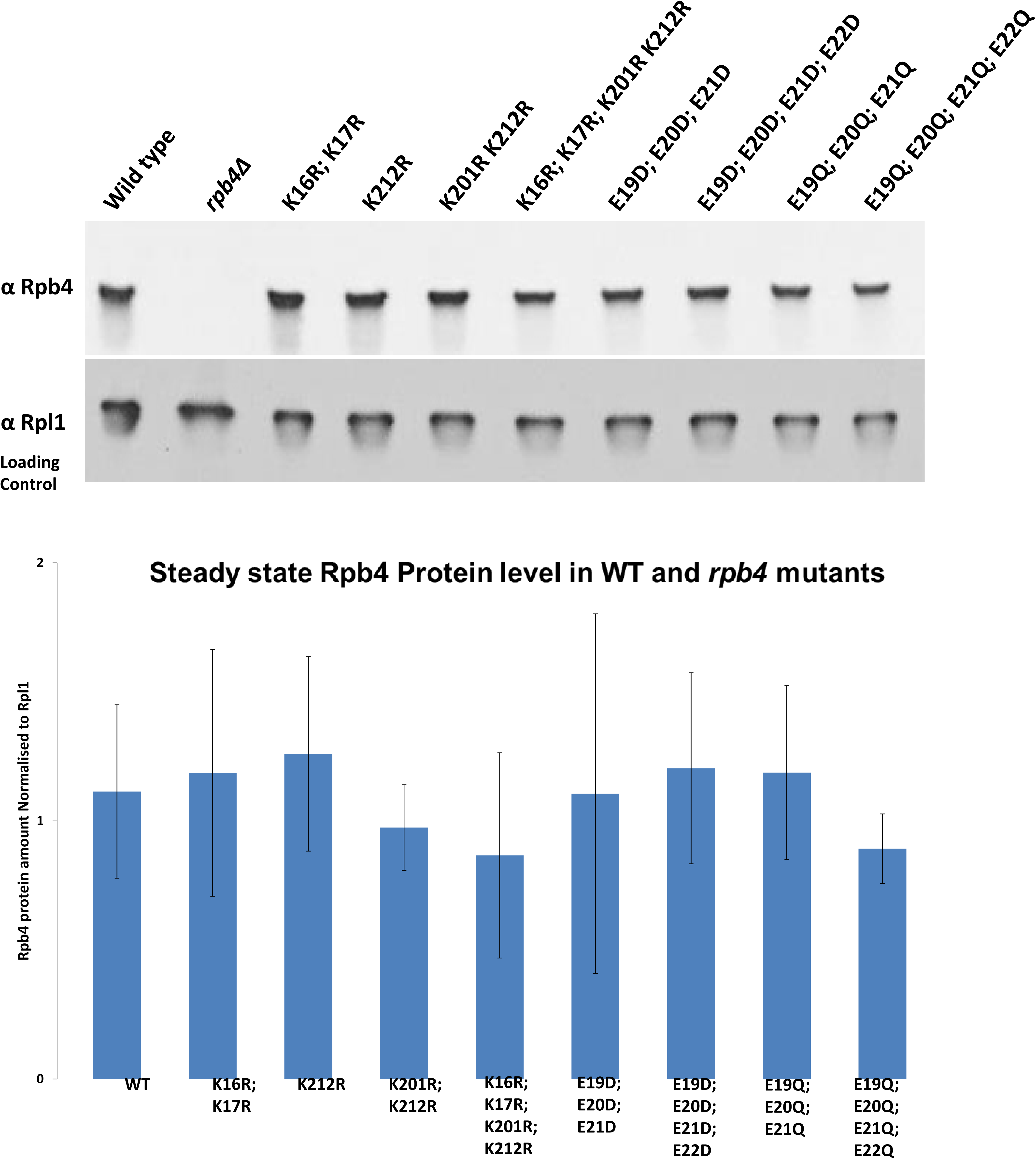
The steady state protein level of our Rpb4 mutant forms is comparable to that of WT Rpb4. (A) Whole cell extracts of WT and indicated *rpb4* mutants were prepared, and equal amount of total protein (50 µg) was resolved on a polyacrylamide gel followed by western analysis with anti-Rpb4 antibodies, and with anti-RplA protein, which served as the loading control. The protein band intensities were quantified using ImageQuant LAS 4010 Imaging & detection system (GE Healthcare Life sciences). The Rpb4 protein level was normalized to Rpl1 protein level for each strain. (B) The average of normalized Rpb4 levels from three biological replicates is plotted in this histogram. Error bars represent standard deviation from the three replicates.

**Supplemental Fig. 5. Related to Fig. 6.**
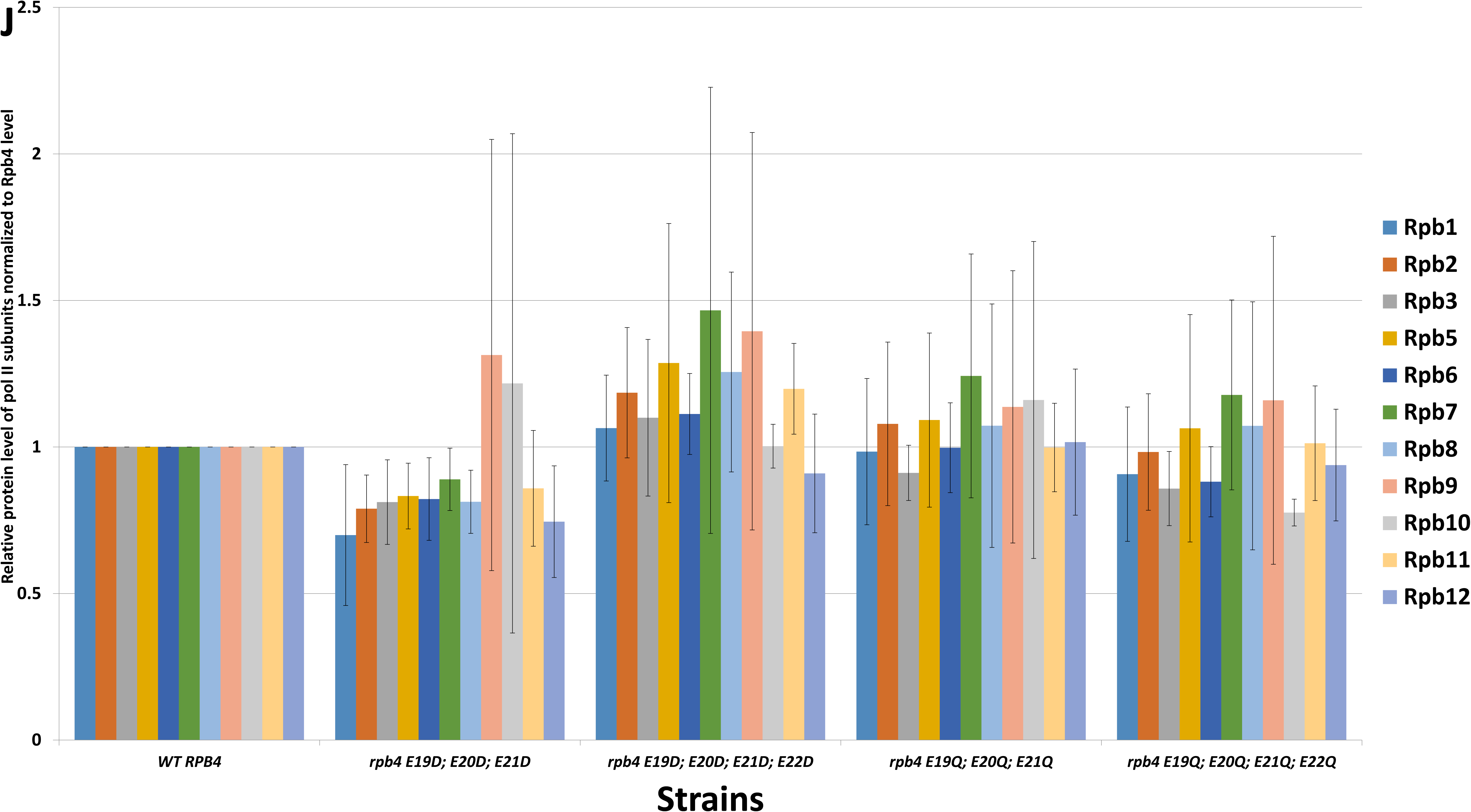
Rpb4 mutant forms bind Pol II comparably to WT. Affinity-purified Rpb4 samples, obtained from the indicated strains, were analyzed by mass spectrometry as in Fig. 6. The protein level of each subunit in each mutant was normalized to its corresponding protein in WT that was defined arbitrarily as 1. Error bars represent SD of three biological replicates.

**Supplemental Fig. 6. Related to Fig. 7.**
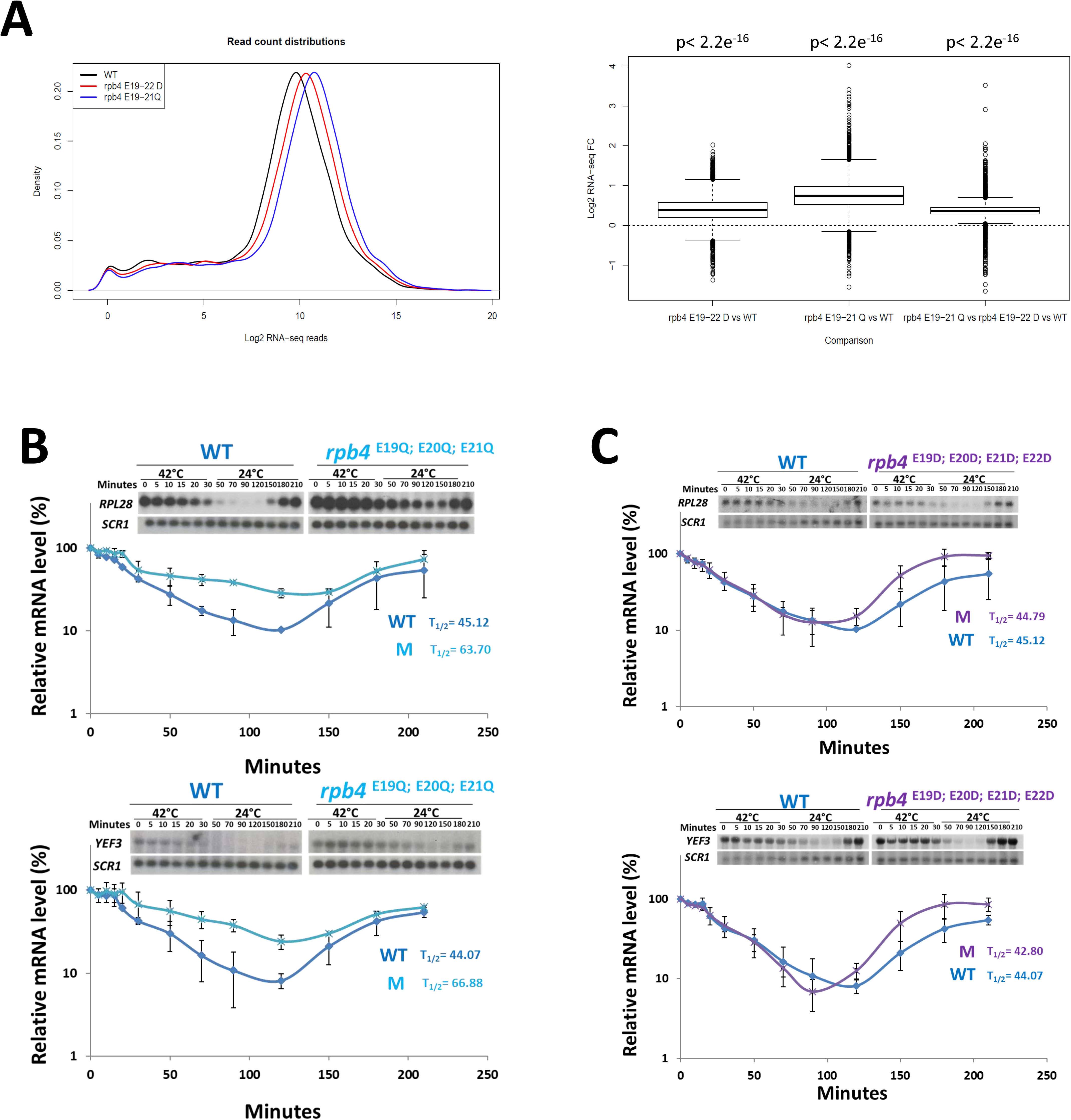
Mutations in Rpb4 E19-22 motif affect mRNA buffering. (A) RNA-seq was performed as described in Fig. 7. Left panel - a density plot of the average normalized reads in each gene for each strain. Right panel – a box plot representing the log2 fold change for each gene between different conditions. P-values were obtained using the Wilcoxon signed-rank test. To get these values we averaged across replicates and then applied the test to the respective pairs of averaged profiles which are shown in the left panel. (B-C) Was performed as in Fig. 7 C and D, except that different probes were used, as indicated.

